# A novel TaNF-YC10-TaNF-YB1-TabHLH95 module coordinates starch biosynthesis in wheat endosperm

**DOI:** 10.64898/2026.02.18.706539

**Authors:** Yunchuan Liu, Yaojia Wang, Haixia Liu, Xiaolu Wang, David Seung, Tian Li, Hongxia Liu, Jian Hou, Xu Liu, Xueyong Zhang, Chenyang Hao

**Author notes:** These authors contributed equally to this article (Yunchuan Liu, Yaojia Wang). Correspondence: Xu Liu, Xueyong Zhang and Chenyang Hao.

## Abstract

Wheat grain weight and flour quality largely depend on starch biosynthesis, yet the mechanisms by which transcription factors coordinate this process remain poorly understood. In this study, using an integrative strategy that combines genome-wide association analysis with yeast two-hybrid library screening, we identify *TaNF-YC10*, a Nuclear Factor Y transcription factor, as a positive regulator of starch accumulation in the wheat endosperm. Loss of *TaNF-YC10* reduces starch content and alters starch granule size distribution, whereas overexpression enhances starch accumulation and increases grain weight. TaNF-YC10 binds and activates core starch biosynthetic-related genes, including *AGPL1*, *GBSS1*, *YUC11*, and *NF-YB7*, and forms higher-order transcriptional complexes with TaNF-YB1 and TabHLH95 to coordinate multiple regulatory pathways. *TaNF-YC10-A1*-*Hap2* is associated with higher starch content and thousand grain weight and has been selected during wheat breeding in China. Collectively, our findings establish *TaNF-YC10* as a pivotal transcriptional hub in starch regulation and highlight its potential as a target for genetic improvement of grain yield in wheat.

## Introduction

Starch, the major storage carbohydrate in the wheat endosperm, accounts for more than 70% of its dry weight and serves as an important determinant of both grain weight and flour quality (Shevkani et al., 2017). In the developing endosperm, starch biosynthesis is governed by a complex and tightly coordinated network of enzymes, sugar transporters, and regulatory factors that collectively determine starch quantity and composition (Huang et al., 2021). Over the past decades, substantial progress has been made in elucidating the enzymatic machinery and sugar transport process that drives starch synthesis, enabling targeted manipulation of starch quantity and structure (Cakir et al., 2015; Li et al., 2021b; Qiu et al., 2025; Wang et al., 2025).

Alongside these advances, accumulating evidence indicates that transcriptional regulation constitutes an indispensable layer of control, integrating developmental and environmental cues to fine-tune starch synthesis during grain filling (Huang et al., 2021). Studies over the past decade have identified diverse transcription factors (TFs) and chromatin regulators that modulate starch biosynthetic pathways, revealing a multilayered and interconnected regulatory landscape (Liu et al., 2022). For instance, TabZIP28, TaPIL1, TaMYB44 and TaMYB1 were identified as direct regulators of starch synthesis-related genes, modulating starch accumulation and structure (Song et al., 2020; Li et al., 2025a; Liu et al., 2025c; Meng et al., 2025). Grain-specific NAC TFs, NAC019, and NAC100 coordinate the synthesis of starch and seed storage proteins, highlighting a tight coordination between carbon and nitrogen metabolism (Gao et al., 2021; Li et al., 2021a). In addition to these, other TFs such as TaDL, TaB3, TaMADS29, and TaNF-YB1 function cooperatively to regulate starch biosynthesis (Liu et al., 2023a; Liu et al., 2025a). Epigenetic regulators such as the CAMTA2-GCN5 complex enhance *TaSus2* and *TaSBEIc* expression by directly binding to their promoters and increasing H3K9ac and H3K14ac levels (Zhang et al., 2024). Nevertheless, gene regulatory network analyses suggest that numerous TFs remain uncharacterized and may occupy central positions within the starch biosynthetic network (He et al., 2024; Zhao et al., 2024), underscoring that current regulatory models are incomplete.

Comparative analyses across cereals suggest that orthologous transcription factors may provide valuable insights into the transcriptional regulation of starch biosynthesis in wheat (Fu and Xue, 2010; Liu et al., 2016). However, divergence between species and the polyploid nature of the wheat genome limit the direct transfer of functional information. For instance, while NF-YB1 regulates starch biosynthesis in both wheat and rice, its downstream target networks are only partially conserved between the two species (Bai et al., 2016; Feng et al., 2022; Liu et al., 2023a; Liu et al., 2025a). Similarly, promoter variation underlies the functional divergence of ABI19 between wheat and maize (Yang et al., 2020; Liu et al., 2023b). Such divergence highlights the necessity for species-specific dissection of regulatory mechanisms.

Among the TF families implicated in starch regulation, the Nuclear Factor Y (NF-Y) family represents a key class of heterotrimeric complexes composed of NF-YA, NF-YB, and NF-YC subunits (Mantovani, 1999). NF-Y complexes have broad roles in plant growth, development, and metabolism, and accumulating evidence indicates that they also play pivotal roles in endosperm starch biosynthesis (Wu et al., 2025). In rice, NF-YB1 interacts with NF-YC12, MADS14, and bHLH144 to cooperatively regulate starch biosynthesis-related genes, including *AGPL2, Waxy, and YUC11* (Bello et al., 2019; Feng et al., 2022; Xu et al., 2021). OsNF-YC8, 9, and 10 promote amylopectin synthesis by activating *SBE3* (Chen et al., 2025). Furthermore, a NF-Y complex fine-tunes *QT12* expression to coordinate rice grain yield and quality, ultimately contributing to synergistic thermotolerance (Li et al., 2025b). In wheat, NF-YB1 similarly regulates core starch biosynthetic genes, including *AGPL2*, *Sus2* and *ISA1*, and forms transcriptional complexes with bHLH95, B3, and DL14 to fine-tune starch synthesis (Liu et al., 2023c; Liu et al., 2025a). In addition, TaNF-YB7, together with NF-YC6 and NF-YA3 subunits, contributes to the regulation of starch biosynthesis by integrating transcriptional and chromatin-based regulation (Chen et al., 2024b). Nevertheless, it remains unclear how specific NF-YC components function as regulatory hubs to coordinate starch biosynthetic gene expression and protein–protein interaction networks in wheat endosperm.

Here, by integrating genome-wide association studies with protein-protein interaction screening, we identify *TaNF-YC10* as a previously uncharacterized factor involved in starch accumulation in wheat. We show that *TaNF-YC10* directly activates starch synthesis-related genes, including *TaAGPL1*, *TaGBSS1*, *TaYUC11*, and *TaNF-YB7*, and interacts with *TabHLH95* and *TaNF-YB1* to enhance transcriptional activity, positioning it as a regulatory hub within the starch biosynthesis network. Notably, natural variation at the *TaNF-YC10-A1* modulates its regulatory capacity, as we identified a haplotype (*Hap2*) that strongly associated with high starch content and thousand grain weight. Together, these findings identify *TaNF-YC10* as a pivotal transcriptional regulator of starch accumulation and reveal its contribution to natural variation in wheat grain yield.

## Results

### GWAS and Y2H library mapping identify *TaNF-YC10* associated with starch content

To identify genetic factors underlying starch accumulation in the wheat endosperm, we performed a genome-wide association study (GWAS) for total starch content using a linear mixed model (LMM) in a natural panel of 145 resequencing wheat accessions (Hao et al., 2020). Three loci on chromosome 7A exceeded the suggestive threshold (*P* < 1.0 × 10^−5^) (Figure 1A and 1B). We focused on the strongest association signal, which localized to a 496.664 Mb-497.762 Mb linkage disequilibrium (LD) block (Figure 1C and Supplemental Table 1). This region harbours nine high-confidence annotated genes, among which *TraesCS7A02G336700* displayed grain-specific expression (Figure 1D; Borrill et al., 2016). Phylogenetic analysis identified it as *TaNF-YC10-A1* (Supplemental Figure 1A).

**Figure 1.**
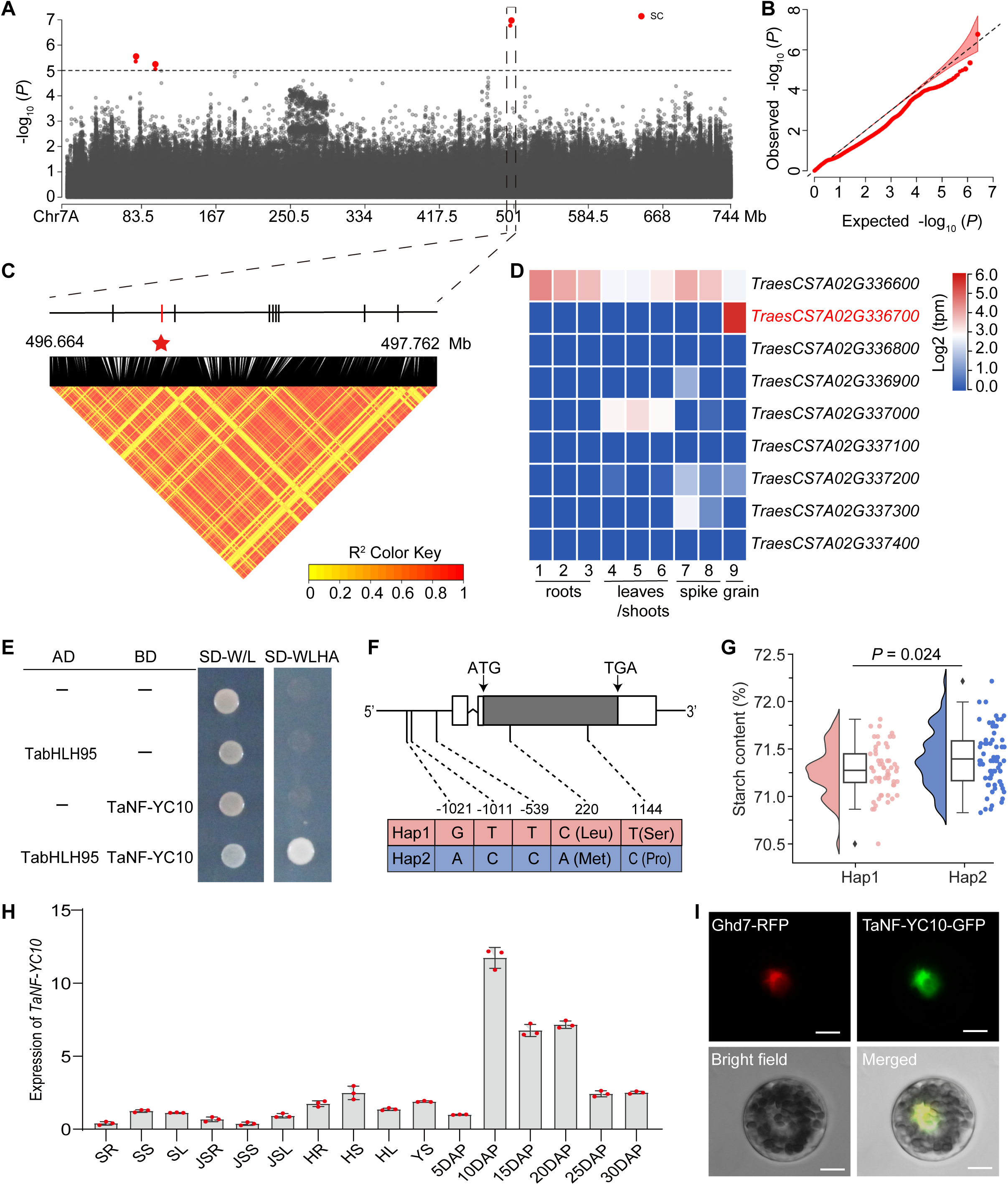
GWAS and Y2H identifies *TaNF-YC10* as a candidate gene regulating starch content in wheat. **(A)** Manhattan plot for total starch content on chromosome 7A. The horizontal dashed line represents the genome-wide suggestive significance threshold (*P* = 1.0×10-5). **(B)** Quantile-quantile plot assessing the performance of the mixed linear model in GWAS. **(C)** Linkage disequilibrium (LD) heatmap surrounding the association peak for total starch content. The black dashed lines indicate the candidate genomic region. Candidate genes are shown as short vertical lines, with *TaNF-YC10-A1* highlighted in red. **(D)** Expression pattern of candidate genes. Data sourced from the Wheat Expression Database (www.wheat-expression.com). **(E)** Y2H confirming the interaction between TaNF-YC10 and TabHLH95. AD, activation domain; BD, binding domain. -LWHA indicates medium lacking Trp, Leu, His, and Ade. **(F)** Allelic variation in the coding sequence (CDS) and 1.8 kb promoter of *TaNF-YC10-A1*. **(G)** Wheat accessions harboring *TaNF-YC10-A1-Hap2* exhibited higher total starch content compared to those with *Hap1*. Data were analyzed using a two-tailed Student’s *t* test (\*\**P* < 0.01). **(H)** Expression pattern of *TaNF-YC10*. Tissues sampled include leaves, stems and roots at seedling stage (SL, SS, and SR), jointing stage (JSL, JSS, and JSR), and heading stage (HL, HS, and HR), 2-cm young spikes (YS), and developing grains at 5, 10, 15, 20, 25, and 30 days after pollination (DAP). Three biological replicates for each experiment with *TaActin* as an internal control. Data are presented as means ± S.D. **(I)** TaNF-YC10 located in the nucleus via co-location with TF Ghd-7 in wheat protoplasts, scale bar = 10 μm.

Yeast two-hybrid (Y2H) library screening indicated that TaNF-YC10 interacts with TabHLH95, a previously identified positive regulator of starch biosynthesis (Liu et al., 2023c). Y2H assays further confirmed the interaction between TaNF-YC10 and TabHLH95 (Figure 1E), implying that TaNF-YC10 may function in the starch synthesis pathway.

Sequence analysis identified five single-nucleotide polymorphisms (SNPs) within the promoter and coding regions of *TaNF-YC10-A1*, which partitioned the natural population into two major haplotypes (Figure 1F). Wheat accessions carrying *Hap2* exhibited significantly higher starch content than those carrying *Hap1* (Figure 1G), indicating functional differentiation at this gene. Expression profiling revealed that *TaNF-YC10* is highly expressed in developing grains, with peaking at 10 days after pollination (Figure 1H). The TaNF-YC10 protein localized to the nucleus in wheat protoplasts and exhibited transcriptional self-activation activity (Figure 1I and Supplemental Figures 1C and Figure 2). Together, these results identify *TaNF-YC10* as a previously uncharacterized regulatory factor associated with starch biosynthesis in wheat.

**Figure 2.**
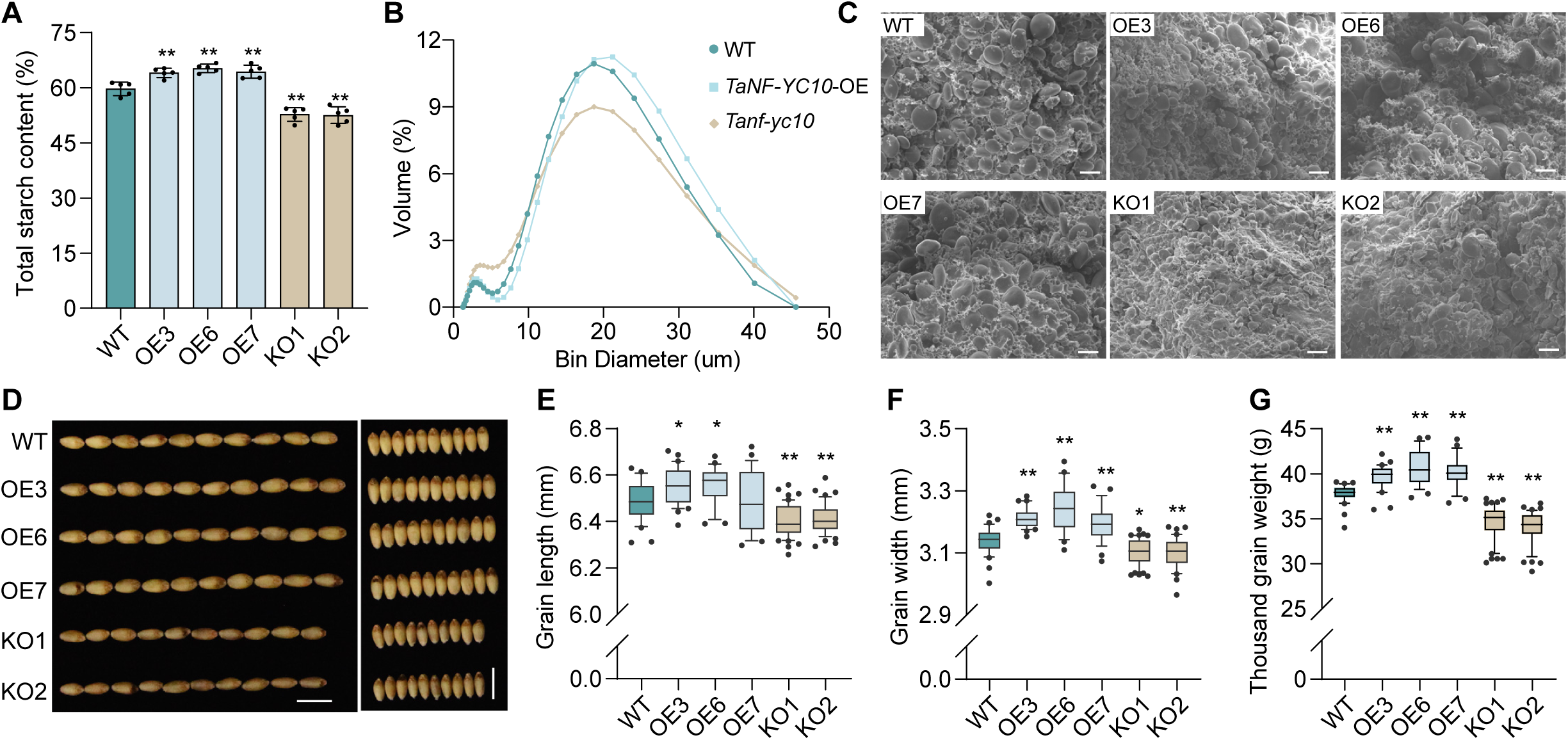
*TaNF-YC10* positively regulates starch content and grain weight. **(A)** Total starch content in WT and *TaNF-YC10* transgenic lines (KO and OE). Each line has five biological replicates. (**B)** Starch granules size distribution of WT and *TaNF-YC10* transgenic lines. **(C)** Scanning electron microscopy of the mature endosperm from WT, *TaNF-YC10* -OE, and -KO lines. n = 6 biological replicates. Scale bars, 20 μm. **(D)** Representative grains from WT and *TaNF-YC10* transgenic lines. Scale bar, 1 cm. **(E-G)** Grain length (**E**), grain width (**F**), and thousand grain weight (**G**) of WT and *TaNF-YC10* transgenic lines. Each line included at least 25 individual plants. Data were analyzed using one-way ANOVA. \**P* < 0.05, \*\**P* < 0.01.

### TaNF-YC10 positively regulates starch content and grain weight in wheat

To investigate the biological function of *TaNF-YC10*, we generated both overexpression (OE) and knockout (KO) lines in the wheat cultivar Fielder. Two independent KO lines carrying large deletions and three OE lines exhibiting significantly elevated *TaNF-YC10* expression were selected for phenotypic analyses (Supplemental Figure 3A and 3B).

Assessment of mature grains showed that total starch content was significantly increased in *TaNF-YC10*-OE lines but markedly reduced in KO lines compared to the wild type (WT), with consistent phenotypes across two growing seasons (Figure 2A and Supplemental Figure 3C). Consistently, OE line exhibited a higher proportion of large starch granules than the WT, whereas KO grains had a higher proportion of small granules (Figure 2B). These trends were further confirmed by scanning electron microscopy, which revealed enlarged starch granules in OE lines and reduced granule size in KO lines (Figure 2C). In addition, amylose content was significantly increased in OE lines and decreased in KO lines (Supplemental Figure 3E). Collectively, these findings demonstrate that *TaNF-YC10* is a positive regulator of starch biosynthesis in wheat.

To assess the agronomic effects of *TaNF-YC10*, we evaluated grain traits of the transgenic lines under field conditions. Compared with the WT, OE lines consistently produced larger grains (Figure 2D). Grain length was significantly increased in OE3 and OE6 during the 2023 spring growing season (Figure 2E), and in OE6 and OE7 during the 2023 winter growing season (Supplemental Figure 3F). Notably, grain width was significantly increased in all three OE lines across both growing conditions (Figure 2F and Supplemental Figure 3G). Consistent with these changes, OE lines showed increased thousand grain weight (TGW), whereas KO lines showed the opposite trend (Figure 2G and Supplemental Figure 3H). The recurrence of these phenotypes across two growing seasons supports a stable positive contribution of *TaNF-YC10* to grain weight.

Additional agronomic traits were evaluated during the 2023 winter growing season. Spike length showed a modest but significantly increased in both OE and KO lines (Supplemental Figure 3L). Other agronomic traits, including plant height, spikelet number, grain number per spike, and tiller number, showed either no significant differences from the WT or inconsistent changes among transgenic lines, with a unclear trend with *TaNF-YC10* expression levels (Supplemental Figure 3G-3N).

### TaNF-YC10 directly regulates starch biosynthesis-related transcriptional programs

To investigate the molecular basis of *TaNF-YC10*-dependent regulation of starch biosynthesis, we performed transcriptome profiling of developing grains at 12 days after pollination (DAP) from *TaNF-YC10* KO and WT plants. In total, 5,319 genes were differentially expressed, among which 2,643 were significantly downregulated in the KO lines (Figure 3A and Supplementary Data 1). Gene Ontology (GO) enrichment analysis revealed that these downregulated genes were predominantly associated with starch synthase activity, amyloplast development, small-molecule biosynthesis, chromatin assembly, and transcriptional regulation (Figure 3B and Supplementary Data 1). These results suggest that loss of *TaNF-YC10* leads to broad transcriptional repression of processes closely linked to transcription regulation and starch biosynthesis.

**Figure 3.**
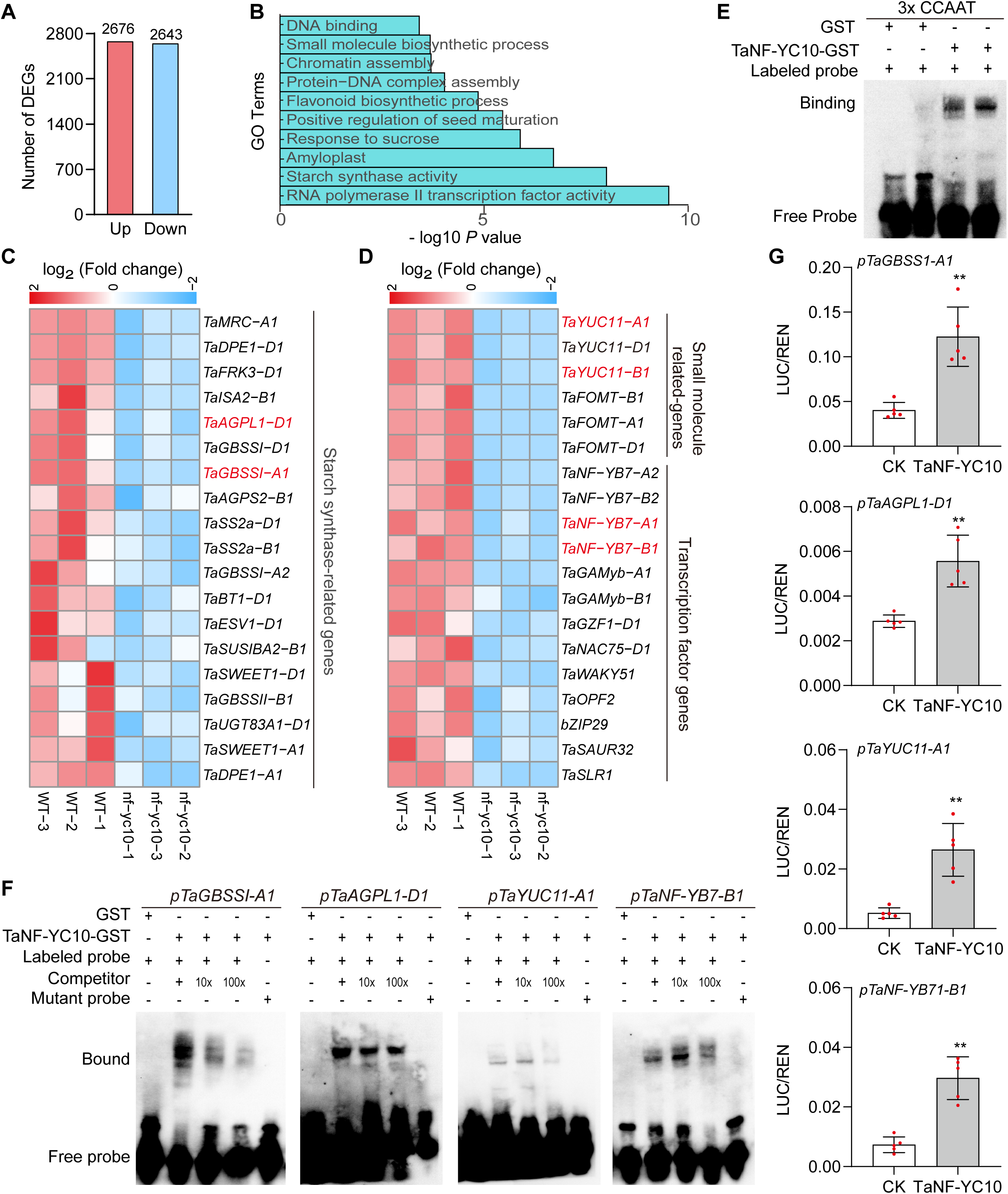
Pathways and downstream genes regulated by *TaNF-YC10*. **(A)** Number of up-regulated and down-regulated genes in 12-DAP *Tanf-yc10* mutants. **(B)** GO enrichment analysis of downregulated DEGs in *Tanf-yc10* mutant. Heatmap of starch biosynthesis-related genes **(C)**, small molecule related genes, and transcription factor genes **(D)** based on RNA-Seq data. **(E)** EMSA showing the direct binding of TaNF-YC10 to CCAAT motif. **(F)** EMSA showing the direct binding of TaNF-YC10 to the promoters of *TaGBSS1-A1*, *TaAGPL1-D1*, *TaYUC11-A1* and *TaNF-YB7-B1*. **(G)** TaNF-YC10 enhanced the transcriptional activity of the *TaGBSS1-A1*, *TaAGPL1-D1*, *TaYUC11-A1* and *TaNF-YB7-B1*. promoters. Statistical significance was determined using a two-tailed Student’s *t* test, \*\**P* < 0.01.

Out of all known starch synthesis genes, most were downregulated in KO lines, including *TaGBSS1*, *TaAGPL1*, *TaBT1*, *TaAGPS2*, *TaISA2*, and *TaSS2a* (Figure 3C). The sugar transport gene *TaSWEET1-A1/D1* and the B-type starch granule regulatory gene *TaMRC-A1* were also decreased expression (Figure 3C; Chen et al., 2024a). Notably, *TaYUC11-A1/B1/D1*, encoding a key enzyme in the indole-3-acetic acid (IAA) biosynthesis pathway and previously implicated in promoting starch accumulation and endosperm development (Xu et al., 2021; Kabir et al., 2021), was markedly downregulated in KO lines (Figure 3D). In addition, numerous TF genes were repressed in the KO background (Figure 3D), including *TaNF-YB7*, which mediates H3K27 methylation and positively regulates starch synthesis (Chen et al., 2024b). By contrast, TFs reported as repressors of starch synthesis, including *NAC019* and *MADS7* (Liu et al., 2020; Zhang et al., 2018), were upregulated in the KO lines (Supplemental Figure 4E). Collectively, these transcriptional changes support the notion that *TaNF-YC10* functions as an upstream regulatory node coordinating transcriptional programs governing starch biosynthesis.

### TaNF-YC10 directly binds CCAAT motifs to activate starch-biosynthetic genes

NF-Y transcription factors typically recognize CCAAT *cis*-elements, and NF-YC subunits have been reported to directly interact with these motifs (Romier et al., 2003; Xiong et al., 2019). Consistent with this, electrophoretic mobility shift assays (EMSA) showed that TaNF-YC10 directly binds to CCAAT elements (Figure 3E). Promoter analysis identified conserved CCAAT motifs in the promoters of key genes involved in starch biosynthesis, IAA synthesis and NF-Y complex formation, including *TaAGPL1-D1*, *TaGBSS1-A1*, *TaYUC11-A1/B1*, and *TaNF-YB7-A1/B1* (Supplemental Figure 4A). EMSA further confirmed direct binding of TaNF-YC10 to the CCAAT-containing promoter fragments of these genes (Figure 3F and Supplemental Figure 4B). Moreover, dual-luciferase reporter (DLR) assays further demonstrated that TaNF-YC10 significantly enhances the transcriptional activity of all above tested promoters (Figure 3G and Supplemental Figure 4C and 4D).

Collectively, these findings indicate that TaNF-YC10 as a direct transcriptional activator of multiple functionally distinct genes central to starch biosynthesis and endosperm development, supporting its role as an upstream regulator that coordinates metabolic, hormonal and epigenetics to promote starch accumulation and grain weight.

### TaNF-YC10 interacts with TabHLH95 to cooperative regulate starch synthesis

We previously identified TabHLH95 as a positive regulator of starch accumulation and a potential interactor of TaNF-YC10 (Liu et al., 2023c). Yeast two-hybrid (Y2H) assays confirmed a physical interaction between TaNF-YC10 and TabHLH95 mediated by the C-terminal region of TaNF-YC10 (Supplemental Figure 5A and 5B). Co-immunoprecipitation (Co-IP) and Luciferase complementation imaging (LCI) assays provided additional evidence that TaNF-YC10 and TabHLH95 associate in *vivo* (Figure 4A and Supplemental Figure 5C). Bimolecular fluorescence complementation (BiFC) showed this interaction is localized the to the nucleus in *N. benthamiana* leaves (Figure 4B).

**Figure 4.**
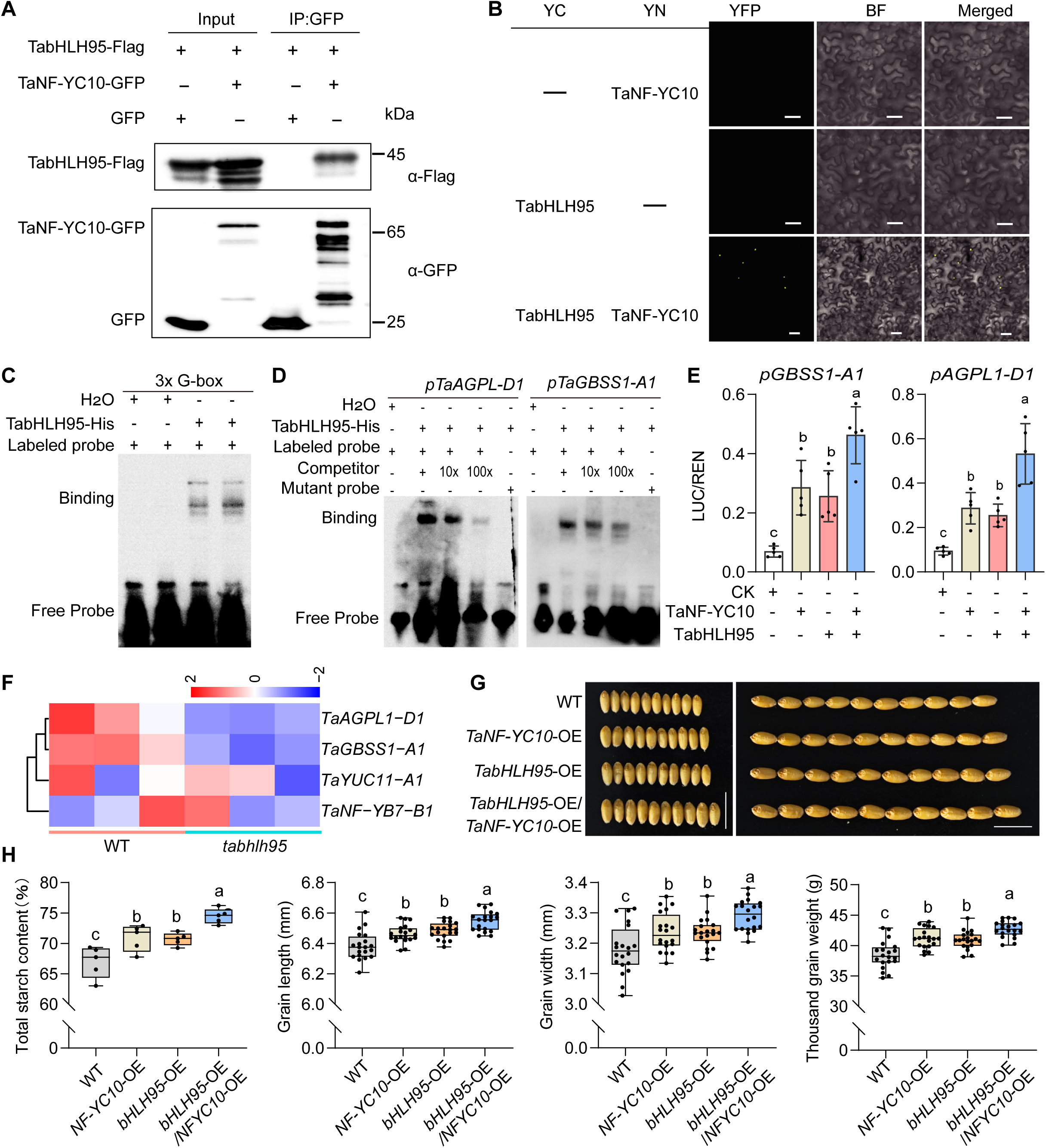
TaNF-YC10 cooperates with TabHLH95 to control starch synthesis and grain weight. **(A)** Co-IP assays showing the interaction between TaNF-YC10 and TabHLH95. **(B)** BiFC assays demonstrating the interaction between TaNF-YC10 and TabHLH95. Scale bar = 10 μm. “-” represents empty carrier. **(C)** EMSA assays demonstrating the binding of TabHLH95 to the G-box. **(D)** EMSA showing the direct binding of TabHLH95 to the promoters of *TaGBSS1-A1*, *TaAGPL1-D1*. **(E)** DLR assays demonstrating that TabHLH95 cooperates with TaNF-YC10 to activate the *TaGBSS1-A1* and *TaAGPL1-D1* promoters. Data represent five biologically independent samples. **(F)** Heatmap of TaNF-YC10 downstream genes based on RNA-seq data from the *tabhlh95* mutant. **(G)** Representative grains in WT, *TaNF-YC10*-OE, *TabHLH95*-OE, and their F_3_ progeny. Scale bar = 1 cm. **(H)** Total starch content (n = 5), grain length (n = 20), grain width (n = 20), and thousand grain weight (n = 20) in WT, *TaNF-YC10*-OE, *TabHLH95*-OE, and their F_3_ progeny. For (**E)** and (**H)**, statistical significance was determined by one-way ANOVA, with different lowercase letters indicating significant differences at *P* < 0.05.

We next examined the functional role of TabHLH95. EMSA assays showed that it binds G-box elements, canonical targets of bHLH transcription factors (Figure 4C; Toledo-Ortiz et al., 2003). G-box motifs were identified in the promoters of *TaAGPL1-D1*, *TaNF-YB7-B1*, *TaYUC11-A1*, and *TaGBSS-A1* (Supplemental Figure 4A). EMSA confirmed direct binding of TabHLH95 to these promoters (Figure 4D and Supplemental Figure 5E). DLR assays revealed that TabHLH95 activated *TaGBSS1-A1* and *TaAGPL1-D1* (Figure 4E), but not *TaYUC11-A1* and *TaNF-YB7-B1* (Supplemental Figure 5F and 5G), consistent with their expression changes in *TabHLH95*-KO lines (Figure 4F and Supplemental Figure 5H and Table 2). Notably, *TabHLH95* co-expression with *TaNF-YC10* further enhanced TaNF-YC10-mediated promoter activation (Figure 4E and Supplemental Figure 5F and 5G), indicating functional synergy between the two factors.

Our previous study demonstrated that *TabHLH95* positively regulates starch biosynthesis and grain weight (Liu et al., 2023c). Here, we re-assessed the effects of *TabHLH95* by re-sowing two previously generated OE lines in field. Consistent with our earlier findings, both *TabHLH95*-OE lines exhibited increased starch content, enlarged grain size, and higher grain weight compared with the WT (Supplementary Figure 5I and 5J).

To assess the genetic interaction between *TaNF-YC10* and *TabHLH95*, we generated double OE lines by crossing method. In the F_3_ generation, grains from the double OE plants exhibited significantly higher starch content than those from the WT and single-gene OE line (Figure 4H). Moreover, grain size and TGW were increased to a greater extent in the double OE plants (Figure 4G and 4H), indicating an additive effect. These results support the notion that *TaNF-YC10* and *TabHLH95* cooperatively enhance starch accumulation and grain weight.

### TaNF-YC10 interacts with TaNF-YB1 to form a transcriptional complex

Our previous study showed that TabHLH95 interacts with TaNF-YB1 and promotes starch biosynthesis (Liu et al., 2023c). Given that TaNF-YC10 also interacts with TabHLH95 and exhibits an additive effect on transcriptional activation, we next examined whether TaNF-YC10 associates with TaNF-YB1.

Using Y2H, LCI, BiFC, and Co-IP assays, we confirmed that TaNF-YC10 physically interacts with TaNF-YB1 both in vitro and in vivo (Figure 5A-D). In addition, EMSA showed that TaNF-YB1 binds to the promoters of *TaGBSS1-A1*, *TaAGPL1-D1*, *TaNF-YB7-A1/-B1*, and *TaYUC11-A1/B1* (Figure 5E and Supplementary Figure 6A). To assess the functional consequence of this interaction, DLR assays revealed that TaNF-YB1 activates these target promoters, and that co-expression of *TaNF-YC10* further enhanced this activation (Figure 5F and 5G, and Supplementary Figure 6B), indicating an additive transcriptional effect. Together, these results support a model in which TaNF-YC10 cooperates with TaNF-YB1 to form a transcriptional activation complex regulating genes involved in starch biosynthesis.

**Figure 5.**
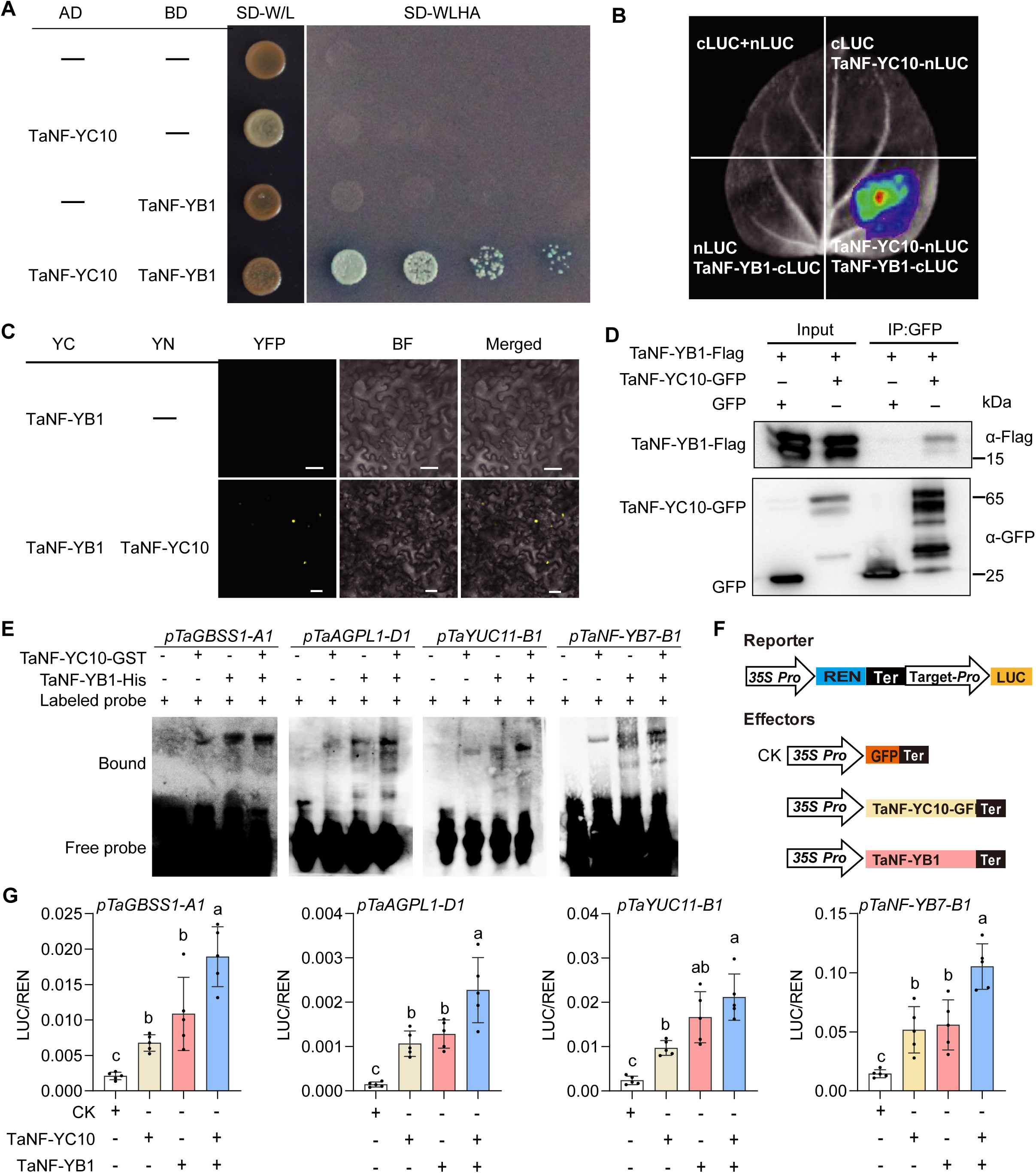
TaNF-YC10 cooperates with TaNF-YB1 to control starch synthesis. **(A)** Y2H assays demonstrating the interaction between TaNF-YC10 and TaNF-YB1. “-” represents empty carrier. **(B)** LCI assays showing the interaction between TaNF-YC10 and TaNF-YB1. **(C)** BiFC assays verifing the interaction between TaNF-YC10 and TaNF-YB1. Scale bar = 10 μm. **(D)** Co-IP assays showing the interaction between TaNF-YC10 and TaNF-YB1. **(E)** EMSA showing the direct binding of TaNF-YC10 and TaNF-YB1 to the promoters of *TaGBSS1-A1*, *TaAGPL1-D1*, *TaYUC11-B1*, and *TaNF-YB7-B1*. **(F)** Schematic representation of reporter and effector constructs used in the DLR assay. LUC, firefly luciferase; 35S, CaMV 35S promoter; REN, Renilla luciferase; Ter, terminator. **(G)** DLR assays demonstrating that TaNF-YB1 cooperates with TaNF-YC10 to activate the *TaGBSS1-A1*, *TaAGPL1-D1*, *TaYUC11-B1*, and *TaNF-YB7-B1* promoters. Data were obtained from five biologically independent samples. Statistical significance was determined by one-way ANOVA, with different lowercase letters indicating significant differences at *P* < 0.05.

### *TaNF-YC10-A1-Hap2* is a favoured in China wheat breeding

To explore the molecular basis underlying the functional divergence between the two *TaNF-YC10-A1* haplotypes, we first analyzed sequence variation in their regulatory and coding regions. The five SNPs within the promoter region did not overlap with conserved cis-elements (Figure 6A), and *TaNF-YC10* transcript abundance was indistinguishable between haplotypes in natural populations (Figure 6B), indicating that differences in transcriptional abundance are unlikely to underlie their functional divergence.

**Figure 6.**
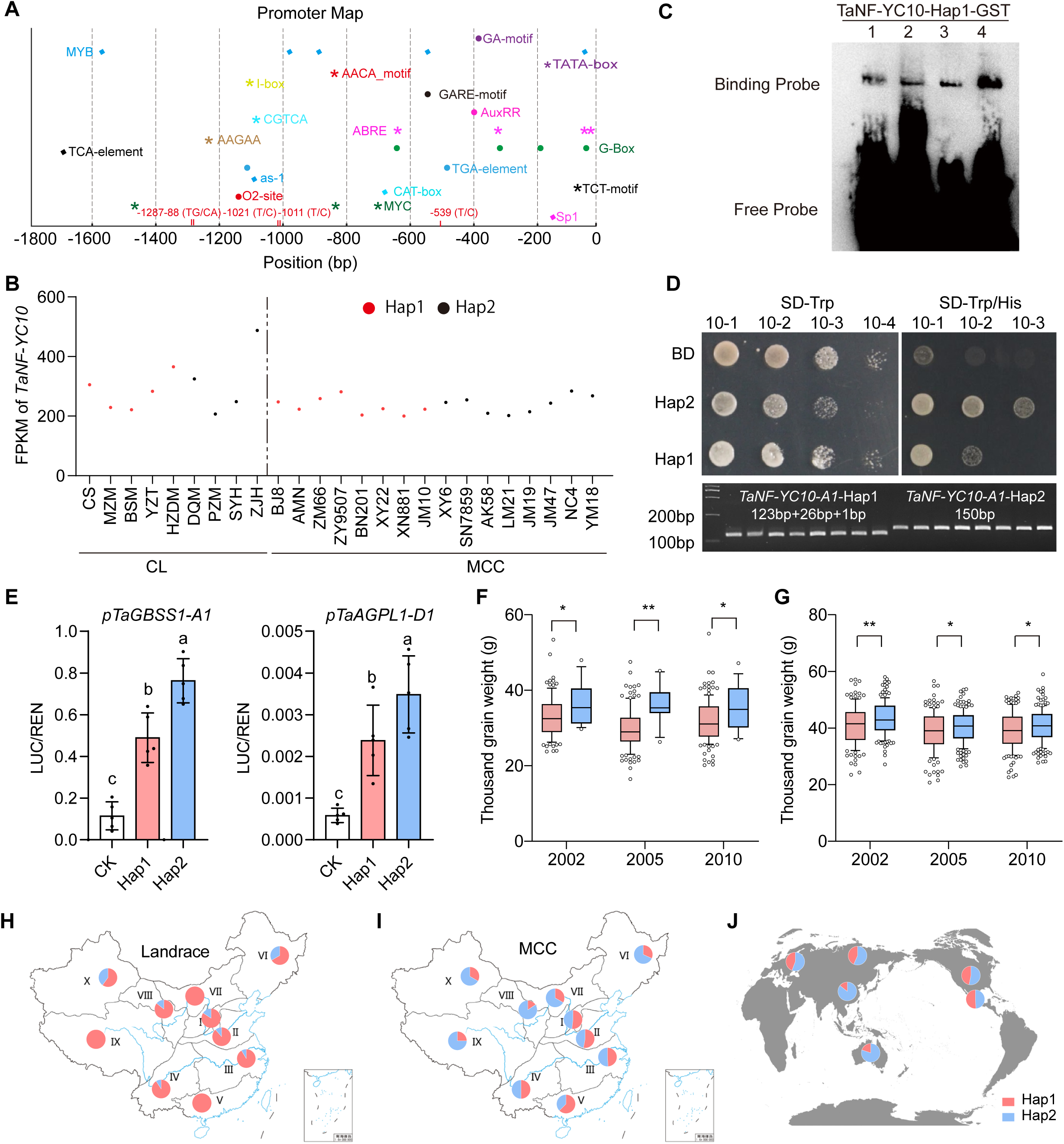
Characterization of *TaNF-YC10-A1* promoter, haplotypes, and functional effects on grain traits. **(A)** Promoter map of *TaNF-YC10-A1* showing cis-elements in the ∼1.8 kb upstream region. Key motifs are indicated at their relative positions (bp) upstream of the start codon. **(B)** Expression levels (FPKM) of *TaNF-YC10-A1* in different wheat assessions. *Hap1* (red) and *Hap2* (black). CL, Chinese landrace; MCC, modern Chinese cultivar. **(C)** EMSA showing binding of TaNF-YC10-A1-Hap1-GST binds to the promoters of *TaAGPL1-D1* (1), *TaGBSS1-A1* (2), *TaYUC11-B1* (3) and *TaNF-YB7-B1* (4). **(D)** Top panel: Yeast assay showing differential transcriptional activity of TaNF-YC10-A1 Hap1 and Hap2. Bottom panel: Development of a dCAPS marker based on the SNP_-1021. **(E)** DLR assays showing transcriptional activation of the *TaGBSS1-A1* and *TaAGPL1-D1* promoters by TaNF-YC10-A1 Hap1 and Hap2. Data represent mean ± SD from five biologically independent samples (one-way ANOVA, lowercase letters represent *P* < 0.05). **(F-G)** Thousand-grain weight of wheat accessions in the landrace (**F**) and modern cultivar (**G**) populations carrying *TaNF-YC10-A1 Hap1* or *Hap2*, measured across three representative years (2002, 2005, 2010). Boxplots indicate the median, interquartile range, and individual data points. In **E-G**, Statistical significance was determined by one-way ANOVA (\**P* < 0.05, \*\**P* < 0.01). **(H-I)** Distribution of *TaNF-YC10-A1* haplotypes in landraces (**H**) and modern cultivars (**I**) across different ecological regions in China. **(J)** Distribution of *TaNF-YC10-A1* haplotypes in different ecological regions globally.

Instead, the two coding-region SNPs were located within the conserved histone-like domain and the C terminus, respectively (Supplemental Figure 7B), and each resulted in an amino acid substitution (Supplemental Figure 7A). EMSA assays showed that the Hap1 protein also displayed binding to the CCAAT motifs present in the promoters of *TaAGPL1-D1*, *TaGBSS1-A1*, *TaYUC11-A1/B1*, and *TaNF-YB7-A1/B1* (Figure 6C and Supplemental Figure 7F), suggesting that these substitutions do not substantially affect DNA-binding affinity.

We next compared the transcriptional activities of the two *TaNF-YC10* haplotype proteins using multiple functional assays. In yeast, *TaNF-YC10*-A1-Hap2 showed stronger transcriptional self-activation than Hap1, a pattern that was also observed in tobacco leaf protoplasts (Figure 6D and Supplementary Figure 7E). Consistently, DLR assays revealed that Hap2 induced higher activation of its downstream gene promoters than Hap1 (Figure 6E and Supplementary Figure 7G). Together, these results suggest that the functional divergence between the two haplotypes primarily reflects differences in intrinsic transcriptional activation capacity.

To evaluate the agronomic significance of this divergence, we developed a dCAPS marker based on SNP_-1021 and genotyped 157 Chinese landraces and 348 modern Chinese cultivars (Figure 6D). Across both natural populations, *Hap2* carriers consistently exhibited significantly higher thousand grain weight than *Hap1* carriers (Figure 6F and 6G). Moreover, across all major agro-ecological zones in China, the frequency of *Hap2* increased markedly from landraces to modern cultivars (Figure 6H and 6I), indicating strong selection for this haplotype during modern breeding. We further examined the global distribution of the two haplotypes in major wheat-growing regions. *Hap2* was present at intermediate to high frequencies worldwide, including Russia (56.6%), Europe (55.7%), North America (53.2%), CIMMYT (50.9%), and Australia (80%) (Figure 6J). Consistent with its phenotypic effects in Chinese populations, these results indicate that *TaNF-YC10-A1-Hap2* is widely distributed in global wheat germplasm. Overall, these findings demonstrate that *TaNF-YC10-A1-Hap2* is a favorable haplotype associated with higher grain weight and has undergone positive selection during wheat breeding in China.

## Discussion

Elucidating the genetic regulation of starch biosynthesis is critical for improving grain weight and end-use quality in wheat. Here, we identify *TaNF-YC10* as a previously uncharacterized transcriptional regulator that coordinates starch accumulation in the endosperm. Although transcriptome analyses had implicated this gene in starch biosynthesis, its molecular function and regulatory mechanisms were unknown (Fang et al., 2022). We further show that TaNF-YC10 enhances starch biosynthesis and grain weight by directly activating key starch biosynthetic-related genes and forming cooperative complexes with TabHLH95 and TaNF-YB1 (Figure 7).

**Figure 7.**
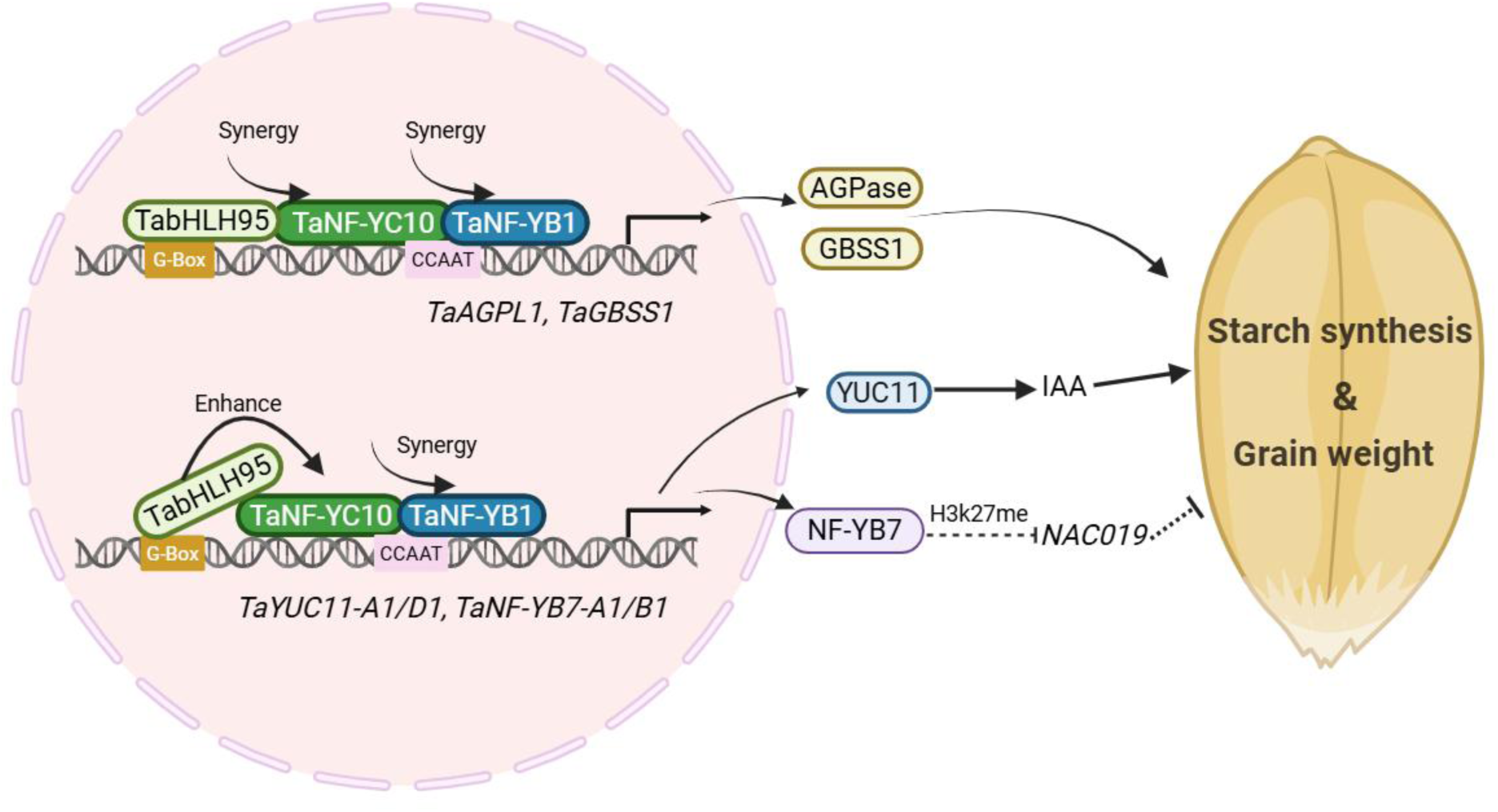
Proposed model illustrating how TaNF-YC10-TaNF-YB1-TabHLH95 *module* regulates starch synthesis and grain weight. TaNF-YC10 interacts with TabHLH95 and TaNF-YB1 to enhance the expression of genes for starch (*TaAGPL1*, *TaGBSS1*) and auxin (*TaYUC11*) biosynthesis, and may also be epigenetically regulated via H3K27me3 (*TaNF-YB7*), promoting starch accumulation and larger grains.

The primary mechanism underlying TaNF-YC10-mediated regulation of starch accumulation involves transcriptional control of core starch biosynthetic pathways. Loss of *TaNF-YC10* is associated with reduced expression of multiple genes involved in starch synthesis, including *BT1*, *AGPS2*, *AGPL1*, *GBSS1, SS2a*, and *SWEET1*, indicating that TaNF-YC10 functions upstream of the starch biosynthetic network (Wang et al., 2019; Shannon et al., 1996; Hogg et al., 2013; James et al., 2003). Within this framework, our results demonstrate that TaNF-YC10 directly activates key starch biosynthetic genes *TaAGPL1-D1* and *TaGBSS1-A1*. *AGPL1* encodes the large subunit of ADP-glucose pyrophosphorylase (AGPase), a rate-limiting enzyme in starch biosynthesis (Ballicora et al., 2004; Jeon et al., 2010; Kang et al., 2013), whereas *GBSS1* is responsible for amylose synthesis (Nakamura et al., 1995; Miura and Sugawara, 1996; Seung, 2020). Reduced expression level of these genes in the *TaNF-YC10*-KO lines provides a mechanistic explanation for the decreased total starch and amylose content observed in mature grains.

In addition to directly regulating core starch biosynthesis genes, *TaNF-YC10* also binds to and activates the promoters of *TaYUC11*-*A1/B1*, linking transcriptional regulation to auxin biosynthesis. Accumulating evidence from cereal crops supports a conserved role for YUC-dependent auxin biosynthesis in endosperm development and starch accumulation (Bernardi et al., 2012; Xu et al., 2021; Ross and McAdam, 2025). Although direct genetic evidence linking *TaYUC11* to starch accumulation is lacking, it is preferentially expressed in developing grains, and its expression correlates with starch synthesis (Kabir et al., 2021). Loss of *TaNF-YC10* markedly reduced *TaYUC11*-*A1/B1/D1* expression, establishing a link between *TaNF-YC10*, auxin, and starch biosynthesis. Together, these findings suggest that TaNF-YC10 may influence starch accumulation by modulating *YUC11*-dependent auxin biosynthesis, consistent with the conserved role of auxin in cereal endosperm. However, direct functional validation of *TaYUC11* in wheat remains to be established.

NF-Y TFs are known to regulate gene expression through epigenetic mechanisms (Oldfield et al., 2019; Hou et al., 2014; Wu et al., 2025). In this study, *TaNF-YC10* may influence starch accumulation through NF-Y associated epigenetic regulation. *TaNF-YB7* forms NF-Y transcriptional complexes that repress downstream targets such as *NAC019* and *Gli*, both directly and indirectly via PRC2-mediated H3K27me3 deposition (Chen et al., 2024a). In *TaNF-YC10* mutants, reduced *TaNF-YB7* expression likely weakens this transcriptional and epigenetic repression, consistent with the observed upregulation of *NAC019*, *Gli*, and *EMF2-like*, accompanied by downregulation of multiple starch biosynthetic genes. Although our results suggest a potential involvement of *TaNF-YC10* in NF-Y modulates H3K27me regulation, genome-wide epigenomic analyses will be essential to further substantiate this regulatory pathway during wheat endosperm development. Collectively, these findings illustrate that TaNF-YC10 functions as an upstream integrator coordinating transcriptional, hormonal, and epigenetic programs to fine-tune starch accumulation.

NF-Y transcription factors regulate plant development in combination with other TFs, thereby expanding their regulatory capacity (Petroni et al., 2012). We find that TaNF-YC10 fine-tunes starch accumulation by interacting with TaNF-YB1 and TabHLH95, analogous to NF-Y-bHLH modules in rice (Liu et al., 2023c, 2025a; Bello et al., 2019). In this study, our results show that each factor exhibits distinct specialization. TaNF-YC10 coordinates metabolic, hormonal, and potential epigenetic pathways. TabHLH95 primarily regulates metabolic genes while enhancing TaNF-YC10-mediated activation of hormonal and epigenetic targets. Their double OE lines demonstrate their synergistic effects, although the dependency of downstream gene regulation remains to be tested. TaNF-YB1 broadly activates core starch and additional grain development related genes and interacts with MADS-box and DROOPING LEAF proteins, similar to rice (Liu et al., 2023a; Liu et al., 2025a, b; Xu et al., 2021), reflecting its role as a versatile integrator. Collectively, these findings reveal a previously unappreciated preference and division of labor among transcription factors, suggesting that transcriptional regulation of starch biosynthesis in wheat endosperm is more complex and coordinated than previously assumed. We propose a model in which TaNF-YC10 serves as a central hub, TaNF-YB1 integrates diverse transcriptional inputs, and TabHLH95 reinforces metabolic regulation to coordinate starch biosynthesis and endosperm development.

The central role of *TaNF-YC10* in the starch regulatory network suggests its potential contribution to grain yield. We found that *TaNF-YC10-A1-Hap2* is associated with higher starch content and increased TGW. Although the causal SNPs have not been genetically confirmed, the two haplotypes show similar expression, suggesting that coding-sequence differences may drive their functional divergence. Hap2 exhibits stronger transcriptional activation, consistent with the C-terminal transactivation domain of TaNF-YC10. The Ser-to-Pro substitution at position 382 may enhance this activity, while the rare Met-to-Leu change at position 74 in Hap1 lies outside the α2 helix and is unlikely to affect NF-Y complex formation or DNA binding (Petroni et al., 2012). This enhanced activation likely explains the superior starch accumulation and grain weight of Hap2. The frequency of *Hap2* is higher in modern cultivars than in landraces, indicating positive selection during wheat breeding. These findings highlight *Hap2* as a favorable allele with potential for improving wheat yield.

In summary, our study identifies TaNF-YC10 as a key transcriptional regulator of starch biosynthesis in the wheat endosperm. TaNF-YC10 functions by interacting with TaNF-YB1 and TabHLH95 to coordinate the expression of core starch biosynthetic genes, thereby influencing grain weight. *TaNF-YC10-A1-Hap2* is associated with higher starch content and grain weight and has been a target of selection during wheat breeding. Together, these findings highlight *TaNF-YC10* as a promising candidate for genetic improvement of starch content and yield in wheat.

## Methods

### Plant materials and growth conditions

For tissue expression analysis, Chinese Spring wheat plants were cultivated in a glasshouse under a 16 h light / 8 h dark photoperiod. For agronomic trait evaluation, WT (Fielder) and *TaNF-YC10* transgenic lines were grown under field conditions in Shunyi, Beijing (116°E, 40°N), during the 2023 spring (early March to mid-July) and winter (mid-October to mid-June) growing seasons. The WT, *TaNF-YC10*-OE, and *TabHLH95*-OE lines, together with the F_3_ generation of *TaNF-YC10*-OE/*TabHLH95*-OE, were grown under field conditions in Shunyi during the 2024 winter growing season. Plants were sown in rows 1.5 m in length, with 30 cm between rows and 10 cm plant spacing. Each genotype was planted in three plots, each comprising four rows. For agronomic trait assessment and harvesting, plants exhibiting uniform growth from the central two rows of each plot were collected. The cultivation conditions of the 145 resequencing wheat accessions used for starch content measurement followed the method previously described (Liu et al., 2025c).

### GWAS for starch content

GWAS for starch content was performed in the 145 re-sequenced land marker cultivars as previously described as previously described (Hao et al., 2020; Liu et al., 2025c). Briefly, genotypic data were analyzed using a mixed linear model implemented in genome wide efficient mixed-model association (GEMMA) software (Zhou and Matthew, 2012), accounting for population structure and kinship. SNPs with *P* < 1 × 10⁻⁵ were considered significantly associated.

### Plasmid construction and plant transformation

The full coding sequence (CDS) of *TaNF-YC10-A1* was cloned into the modified pCAMBIA1305 vector under the control of the *1Bx7* promoter to generate overexpression lines. For CRISPR/Cas9-mediated knockout, two sgRNAs targeting conserved regions of the three homologous were cloned into the pBUE411 vector. *Agrobacterium*-mediated transformation was performed in the wheat cultivar Fielder. Overexpression lines with moderate transcript abundance, as determined by qPCR, were selected for phenotype analyses, while knockout genotypes were confirmed by Sanger sequencing across generations. The *TabHLH95* transgenic plants were generated in a previously study (Liu et al., 2023c).

### Starch phenotype collection

After harvest, mature seeds were oven-dried at 45℃, ground into flour, and passed through a 100 µm sieve. Total starch content was quantified using a commercial starch assay kit (Megazyme) with five biological replicates per genotype. Amylose content was determined using the iodine-binding method with a calibration curve generated from amylose standards (0.4%, 10.6%, 16.2%, 26.5%), also with five biological replicates per genotype. Starch granule size and distribution were analyzed by wet dispersion using a Mastersizer 3000 particle size analyser. In addition, starch granule morphology and size in mature endosperm were examined by scanning electron microscopy (SEM), with three biological replicates per genotype.

### RNA-Seq and qPCR analyses

Total mRNA was extracted from various tissues of Chinese Spring, Fielder and *TaNF-YC10* transgenic lines using TRIzol method, and cDNA was synthesized with the SuperScript IV kit (Invitrogen, USA). qPCR was performed on a LightCycler^®^ 480 (Roche) with 2× SYBR Green Taq (TaKaRa, Japan), with three biological replicates per sample.

Developing grains 12 days post-pollination from Fielder and *TaNF-YC10-*KO lines were collected for RNA-seq. High-quality reads were mapped to the wheat reference genome (IWGSC RefSeq v1.0), and data analysis and visualization were performed using the Novogene Cloud platform.

### Subcellular localization

The CDSs of *TaNF-YC10-A1*, *-B1* and *-D1* were amplified from grain-derived cDNA and cloned into the *35S::*eGFP expression vector. After that, the GFP-fusion constructs were co-transiently expressed with the nuclear marker Ghd7-RFP in wheat protoplasts. After 14 h incubation, fluorescence signals were detected using a laser confocal microscope (LSM980, Carl Zeiss, Germany).

### Yeast-2-hybrid

The CDSs of *TaNF-YB1-A1*, as well as the N- and C-terminal regions of *TaNF-YC10-A1*, were individually cloned into the pGBDT7 vector. The CDS of *TabHLH95-A1* was cloned into the pGADT7 vector. The constructs were co-transformed into the yeast strain AH109 according to a combinatorial strategy. Protein-protein interactions were assessed by monitoring yeast growth on SD/-Trp-Leu-His-Ade medium.

### Co-immunoprecipitation (Co-IP)

To examine the interactions among TaNF-YB1, TaNF-YC10, and TabHLH95, the open reading frames (ORFs) of *TaNF-YB1-A1* and *TabHLH95-A1* were fused to a C-terminal Flag tag, whereas *TaNF-YC10-A1* was cloned into the *35S::*GFP vector. The resulting constructs were transformed into *Agrobacterium tumefaciens* strain GV3101 and co-infiltrated into *Nicotiana benthamiana* leaves. After 48 h, GFP fluorescence was detected using confocal microscopy, and leaf samples were collected for immunoprecipitation and western blot analyses as previously described (Li et al., 2023).

### Luciferase complementation imaging (LCI) assay

For analysis of TaNF-YC10, TaNF-YB1, and TabHLH95 interactions, the CDSs of *TabHLH95-A1* and *TaNF-YB1-A1* were cloned into the cLUC vector, and *TaNF-YC10-A1* was cloned into the nLUC vector. The recombinant constructs were transiently co-expressed *in N. benthamiana* leaves via *A. tumefaciens*-mediated infiltration. After 48 h cultured, leaves were treated with 1mM D-luciferin, and luminescence was recorded to assess protein-protein interactions.

### Bimolecular fluorescence complementation (BiFC)

The ORFs of *TabHLH95-A1* and *TaNF-YB1-A1* were fused to the C-terminus of YFP, while *TaNF-YC10-A1* was cloned into the N-terminal YFP vector. The constructs were transiently co-expressed in *N. benthamiana* leaves via *A. tumefaciens* infiltration. YFP fluorescence was examined 48 hours post-infiltration using a confocal laser scanning microscope (ZEISS LSM 980, Carl Zeiss, Germany).

### Electrophoretic mobility-shift assay (EMSA)

The CDSs of *TaNF-YB1-A1* and *TabHLH95-A1* were fused with 3× His, and *TaNF-YC10-A1-Hap1* and *-2* were inserted into the pGEX4T vector. The recombinant plasmids were introduced into *Escherichia coli* BL21 (DE3), and the His- and GST-tagged proteins were purified using Ni-NTA and glutathione affinity resin, respectively, and concentrated for EMSA. Biotin-labeled DNA probes, corresponding to promoter regions of downstream genes, including the “CCAATA” and “G-box” motifs, were modified at the 3′ end (Probes listed in Supplementary Data 2). EMSA was performed using the LightShift Chemiluminescent EMSA Kit (Thermo Fisher Scientific, 20148) following the manufacturer’s protocol.

### Dual-luciferase reporter assay

The 2 kb promoter regions of downstream genes were cloned into the pGreen-II-0800-Luc vector as reporters (Hellens et al., 2005). The ORFs of *TabHLH95-A1*, *TaNF-YB1-A1*, and *TaNF-YC10-A1* were driven by the *35S* promoter as effectors. Reporter and effector constructs were individually transformed into *A. tumefaciens* GV3101 carrying *p19* gene. The strains (OD_600_ = 1) were mixed at a 1:2 ratio and infiltrated into *N. benthamiana* leaves, with at least of three biological replicates per experiment. LUC and REN activities were quantified using the Dual-Luciferase Reporter Assay System (Promega, USA).

### Data statistics

Statistical analyses were performed in R software (version 4.3.1). Two-group comparisons were conducted using Student’s *t*-test, and differences among multiple groups were assessed by One-way ANOVA.

### Reporting summary

Further information on research design is available in the Nature Research Reporting Summary linked to this article.

## ACKNOWLEDGMENTS

We would like to thank Professor Genying Li (Shandong Academy of Agricultural Sciences) for assistance with wheat transformation. We are also grateful to Dr. Jianhua Huang (John Innes Centre) for his valuable suggestions.

## AUTHOR CONTRIBUTIONS

C.Y.H., X.Y.Z. and X.L. conceived and supervised the project. Y.C.L., Y.J.W. and X.L.W. performed the experiments. H.X.L. conducted the GWAS. Y.J.W., T. L., J.H., and H.X.L. investigated the agronomic phenotypes. Y.C.L. and Y.J.W. wrote the manuscript. D.S., X.Y.Z. and C.Y.H. revised the manuscript.

## SUPPLEMENTARY INFORMATION

**Supplementary Figure 1.** Phylogenetic analysis, domain structure, and transcriptional activity of TaNF-YC10 proteins.

**Supplementary Figure 2.** TaNF-YC10 homoeologous localize to the nucleus in wheat protoplasts.

**Supplementary Figure 3.** Phenotypes of *TaNF-YC10* OE and KO lines.

**Supplementary Figure 4.** TaNF-YC10 directly binds to and activates the promoter of *TaYUC11-B1* and *TaNF-YB7-A1*.

**Supplementary Figure 5.** TabHLH95 interacts with TaNF-YC10 to cooperatively regulate starch biosynthesis and grain weight in wheat.

**Supplementary Figure 6.** TaNF-YC10 cooperates with TaNF-YB1 to activate downstream target genes.

**Supplementary Figure 7.** Natural variation of *TaNF-YC10-A1* leads to differential transcriptional regulation of downstream genes.

**Supplementary Table 1.** Genes in the 7A LD block.

**Supplementary Table 2.** Expression of starch synthesis related genes in *tabhlh95* mutant.

**Supplementary Data 1.** Differentially expressed genes, GO enrichment and data of Hotmap based RNAseq from *Tanf-yc10* mutant vs WT.

**Supplementary Data 2.** List of primers used in this study.

