## Supplemental Table 2 for "A novel TaNF-YC10-TaNF-YB1-TabHLH95 module coordinates starch biosynthesis in wheat endosperm"

**Supplementary Table 2. Expression of starch synthesis related genes in *tabhlh95* mutant.**

| Gene Name | WT-1 | WT-2 | WT-3 | bhlh95-1 | bhlh95-2 | bhlh95-3 |
| --- | --- | --- | --- | --- | --- | --- |
| TaGBSS-A1 | 1.08 | 1.07 | 0.86 | 0.51 | 0.41 | 0.55 |
| TaAGPL-D1 | 1.02 | 1.32 | 0.66 | 0.43 | 0.32 | 0.36 |
| TaYUC11-B1 | 0.83 | 1.18 | 0.99 | 0.79 | 1.04 | 1.06 |
| TaNF-YB7-B1 | 0.97 | 0.92 | 1.11 | 0.95 | 0.92 | 1.09 |
