## Supplemental Table 1 for "A novel TaNF-YC10-TaNF-YB1-TabHLH95 module coordinates starch biosynthesis in wheat endosperm"

| **Chrom** | **Start** | **END** | **Gene02G** | **Strand** | **Function description** |
| --- | --- | --- | --- | --- | --- |
| chr7A | 496.844364Mb | 496.863172Mb | TraesCS7A02G336600 | - | Elongation factor G |
| chr7A | 496.90289Mb | 496.908817Mb | TraesCS7A02G487900LC | + | ubiquitin-protein ligase 4 |
| chr7A | 496.982862Mb | 496.984529Mb | TraesCS7A02G488000LC | + | Vasopressin V1a receptor |
| chr7A | 496.982864Mb | 496.984529Mb | TraesCS7A02G336700 | - | Nuclear transcription factor Y subunit C |
| chr7A | 497.019634Mb | 497.020539Mb | TraesCS7A02G336800 | - | DNA ligase IV-binding protein |
| chr7A | 497.286758Mb | 497.290551Mb | TraesCS7A02G336900 | + | DnaJ subfamily C member 2 |
| chr7A | 497.293389Mb | 497.295659Mb | TraesCS7A02G488100LC | - | F-box domain containing protein |
| chr7A | 497.297026Mb | 497.301506Mb | TraesCS7A02G337000 | + | Phosphoglucosamine mutase family protein |
| chr7A | 497.309563Mb | 497.309892Mb | TraesCS7A02G488200LC | - | Zinc finger BED domain-containing protein |
| chr7A | 497.310282Mb | 497.311225Mb | TraesCS7A02G488300LC | - | BED zinc finger |
| chr7A | 497.311256Mb | 497.312635Mb | TraesCS7A02G337100 | - | Zinc finger BED domain-containing 4 |
| chr7A | 497.313371Mb | 497.316566Mb | TraesCS7A02G337200 | + | Hepatoma-derived growth factor-related protein 2 |
| chr7A | 497.556129Mb | 497.558152Mb | TraesCS7A02G337300 | + | Leucine-rich repeat receptor-like protein kinase |
| chr7A | 497.650003Mb | 497.650399Mb | TraesCS7A02G488400LC | + | RNA-directed DNA polymerase (reverse transcriptase)-related family protein |
| chr7A | 497.650532Mb | 497.658195Mb | TraesCS7A02G337400 | - | Laccase |
| chr7A | 497.692314Mb | 497.69309Mb | TraesCS7A02G488500LC | + | Retrotransposon protein |

**Supplementary Table 1.** **Genes in the 7A LD block**
