## Supplemental Figures for "A novel TaNF-YC10-TaNF-YB1-TabHLH95 module coordinates starch biosynthesis in wheat endosperm"

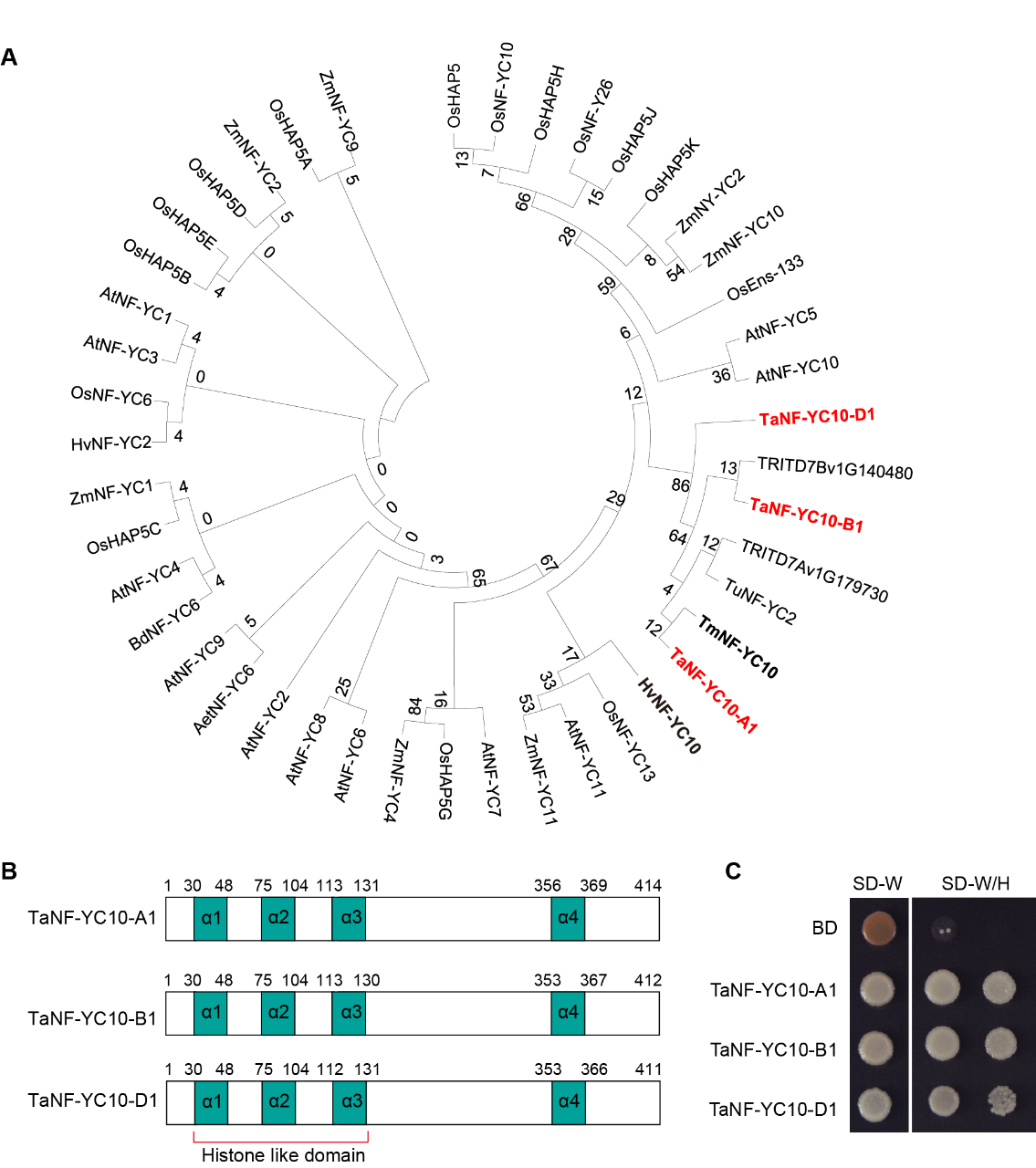

**Supplementary Figure 1. Phylogenetic analysis, domain structure, and transcriptional activity of TaNF-YC10 proteins. (A)** Phylogenetic tree of NF-YC proteins from wheat (*Ta*), rice (*Os*), maize (*Zm*), barley (*Hv*), Arabidopsis (*At*), Brachypodium (*Bd*), and *Triticum urartu* (*Tu*). TaNF-YC10 homologs are highlighted in red. Maximum Likelihood method Bootstrap=1000. The tree was constructed from protein sequences using the maximum likelihood method with 1000 bootstrap repetitions. Numbers on the branches indicate bootstrap values. **(B)** Schematic representation of the protein domain structures of TaNF-YC10-A1, TaNF-YC10-B1, and TaNF-YC10-D1. α-helices (α1–α4) are shown in colored boxes, and the histone-like domain is indicated by a bracket. Numbers above the boxes represent amino acid positions. **(C)** Transcriptional activation activity of TaNF-YC10-A1, TaNF-YC10-B1, and TaNF-YC10-D1 in yeast. Yeast cells transformed with the indicated constructs were grown on SD-Trp (SD-W) or SD-Trp/His (SD-W/H) selective medium. BD, empty vector control.

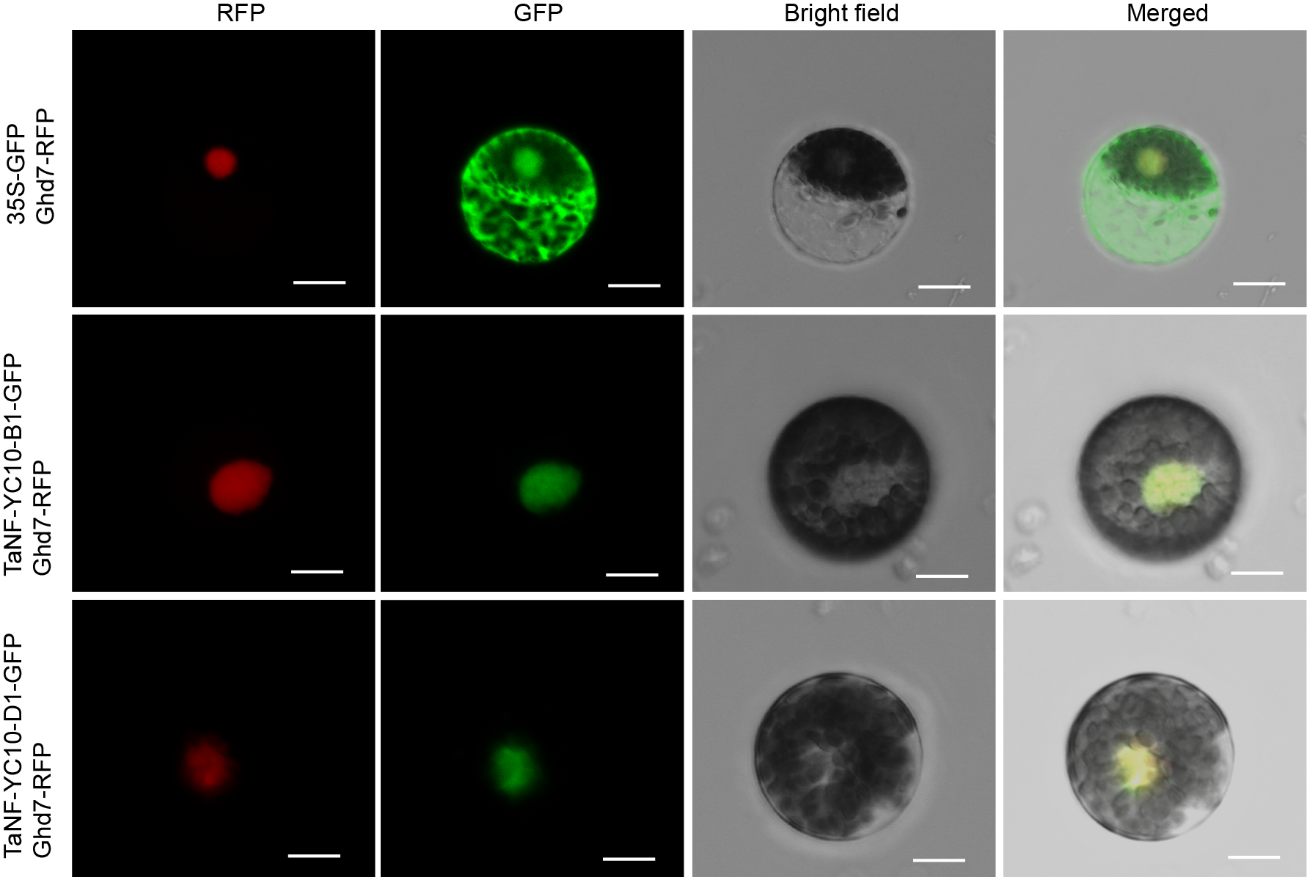

**Supplementary Figure 2. TaNF-YC10 homoeologs localize to the nucleus in wheat protoplasts.** Scale bar = 20 μm.

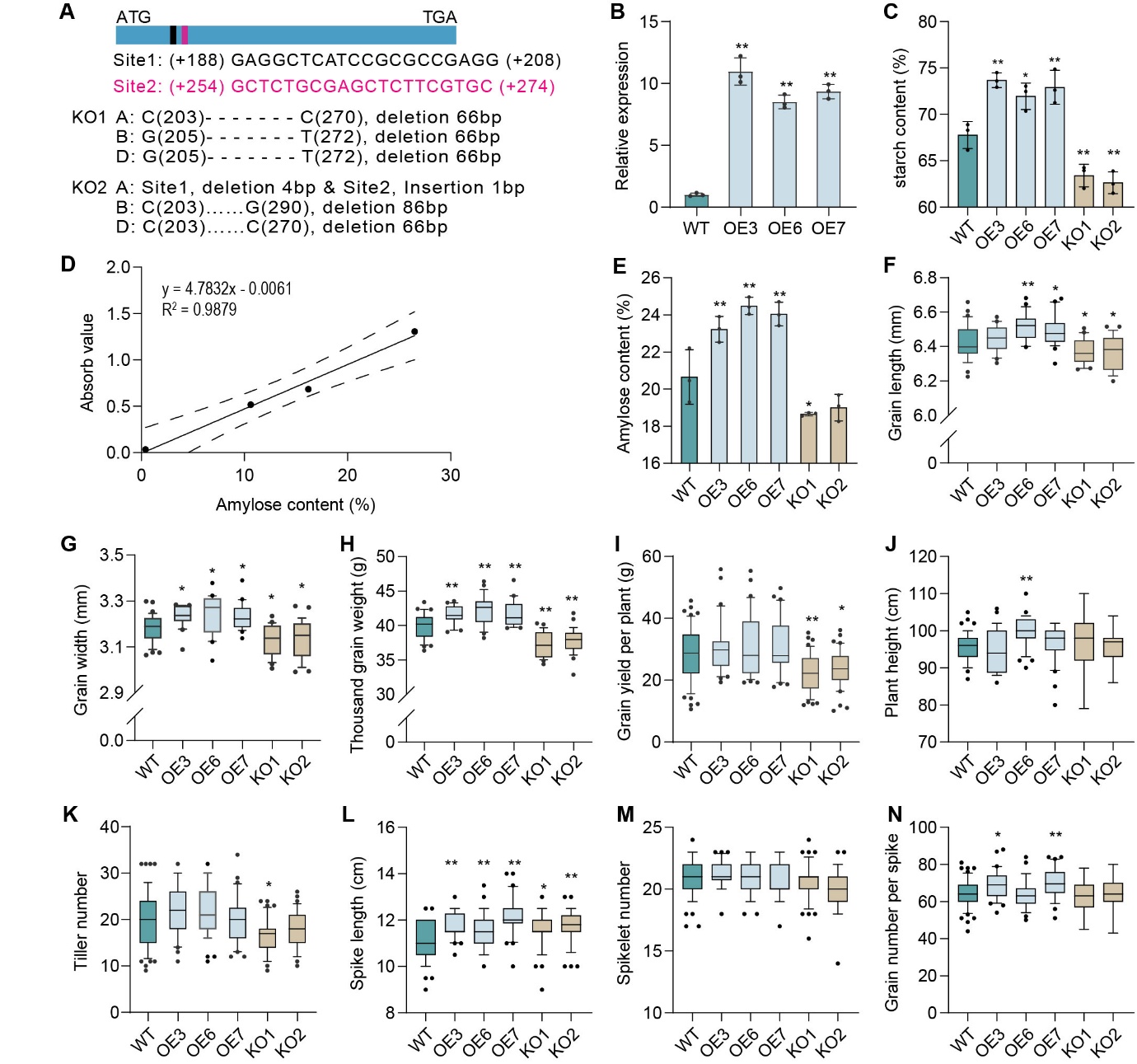

**Supplementary Figure 3. Phenotypes of *TaNF-YC10* OE and KO lines. (A)** The sgRNA sequence used in the CRISPR-Cas9 system for generating *TaNF-YC10* KO lines was designed based on a conserved region shared among the three *TaNF-YC10* homologs. **(B)** Expression levels of *TaNF-YC10* in WT and OE lines. Data represent means ± SD of three biological replicates. *TaActin* was used as an internal control. Statistical significance was determined by one-way ANOVA (**P* < 0.05, ***P* < 0.01). **(C)** Total starch content in WT and *TaNF-YC10* transgenic lines under winter growing condition. **(D)** Standard curve for amylose content based on absorbance. **(E)** Amylose content in WT and *TaNF-YC10* transgenic lines under winter growing condition. For **C** and **E**, data represent means ± SD (n =3). Statistical significance was determined using one-way ANOVA (***P* < 0.01, **P* < 0.05). **(F-N)** Grain length (**F**), grain width (**G**), thousand grain weight (**H**), grain yield per plant (**I**), plant height (**J**), tiller number (**K**), spike length (**L**), spikelet number (**M**), and grain number per spike (**N**) in WT and *TaNF-YC10* transgenic lines under winter growing condition. For **F-N**, data represent means ± SD (n≥15 for agronomic traits). Statistical significance was determined using one-way ANOVA (**P* < 0.05, ***P* < 0.01) compared with WT.

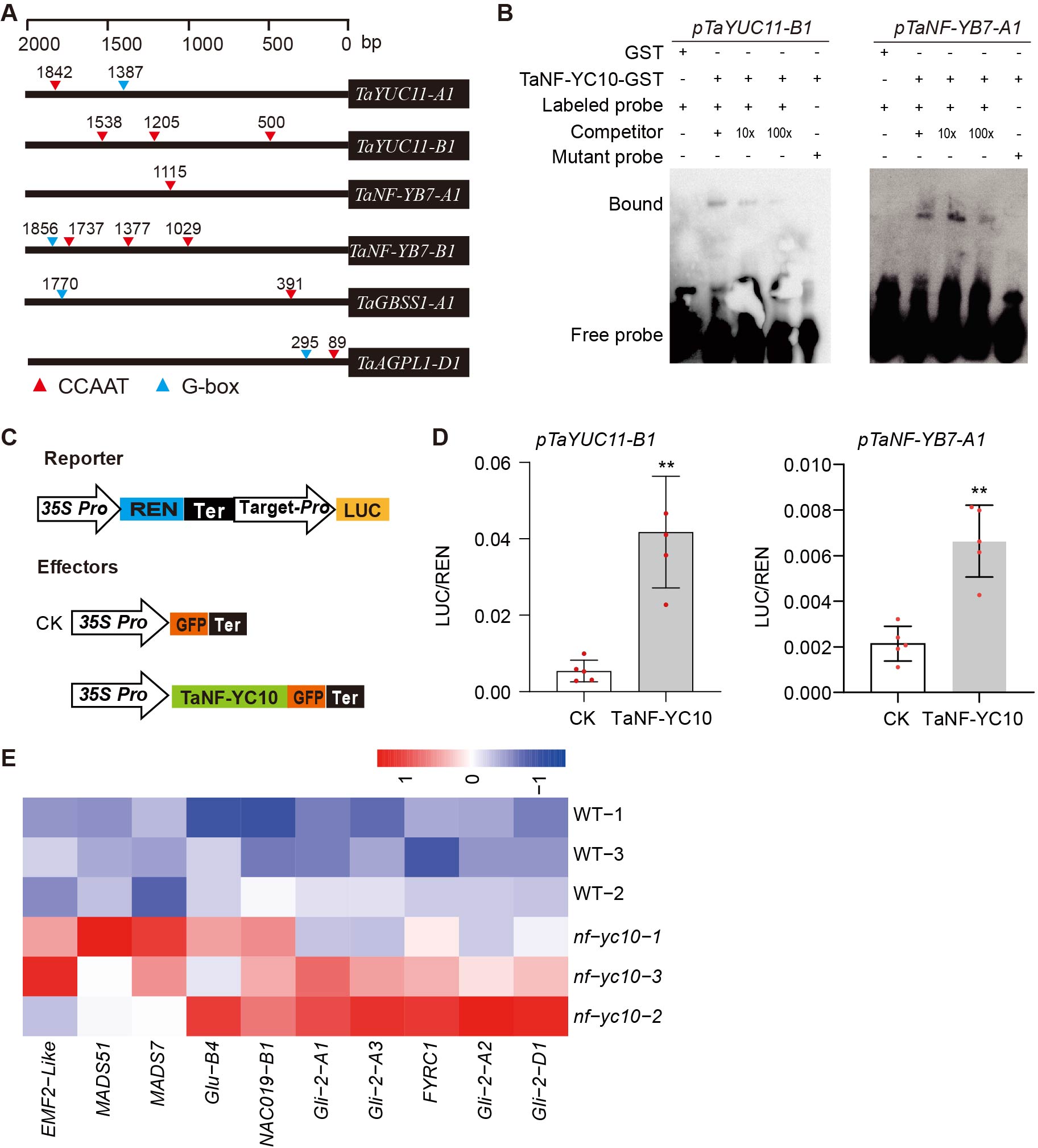

**Supplementary Figure 4. TaNF-YC10 directly binds to and activates the promoter of *TaYUC11-B1* and *TaNF-YB7-A1*. (A)** Schematic representation of promoter regions of *TaYUC11-A1*/*-B1*, *TaNF-YB7-A1*/*-B1*, *TaGBSS1-A1* and *TaAGPL1-D1*. Red triangles indicate CCAAT motifs and blue triangles indicate G-box elements; numbers denote distances (bp) from the translation start site. **(B)** EMSA showing binding of TaNF-YC10 to the promoters of *TaYUC11-B1* and *TaNF-YB7-A1*. GST alone was used as a negative control. Competition assays were performed with 10× and 100× unlabelled probes, and a mutant probe was used to confirm binding specificity. **(C)** Schematic diagram of the dual-luciferase reporter system. Target gene promoters were fused to the LUC reporter, while REN driven by the 35S promoter served as an internal control. Effector constructs expressed GFP (CK) or *TaNF-YC10*-GFP under the *35S* promoter. **(D)** DLR assays showing activation of *TaYUC11-B1* and *TaNF-YB7-A1* promoters by TaNF-YC10 in *Nicotiana benthamiana* leaves. Data are presented as LUC/REN ratios (mean ± SD., n = five biologically independent samples). ***P* < 0.01 (two-tailed Student’s t-test). **(E)** Heatmap showing expression levels of *Gli* and starch synthesis related transcription factor genes in WT and *Tanf-yc10* lines.

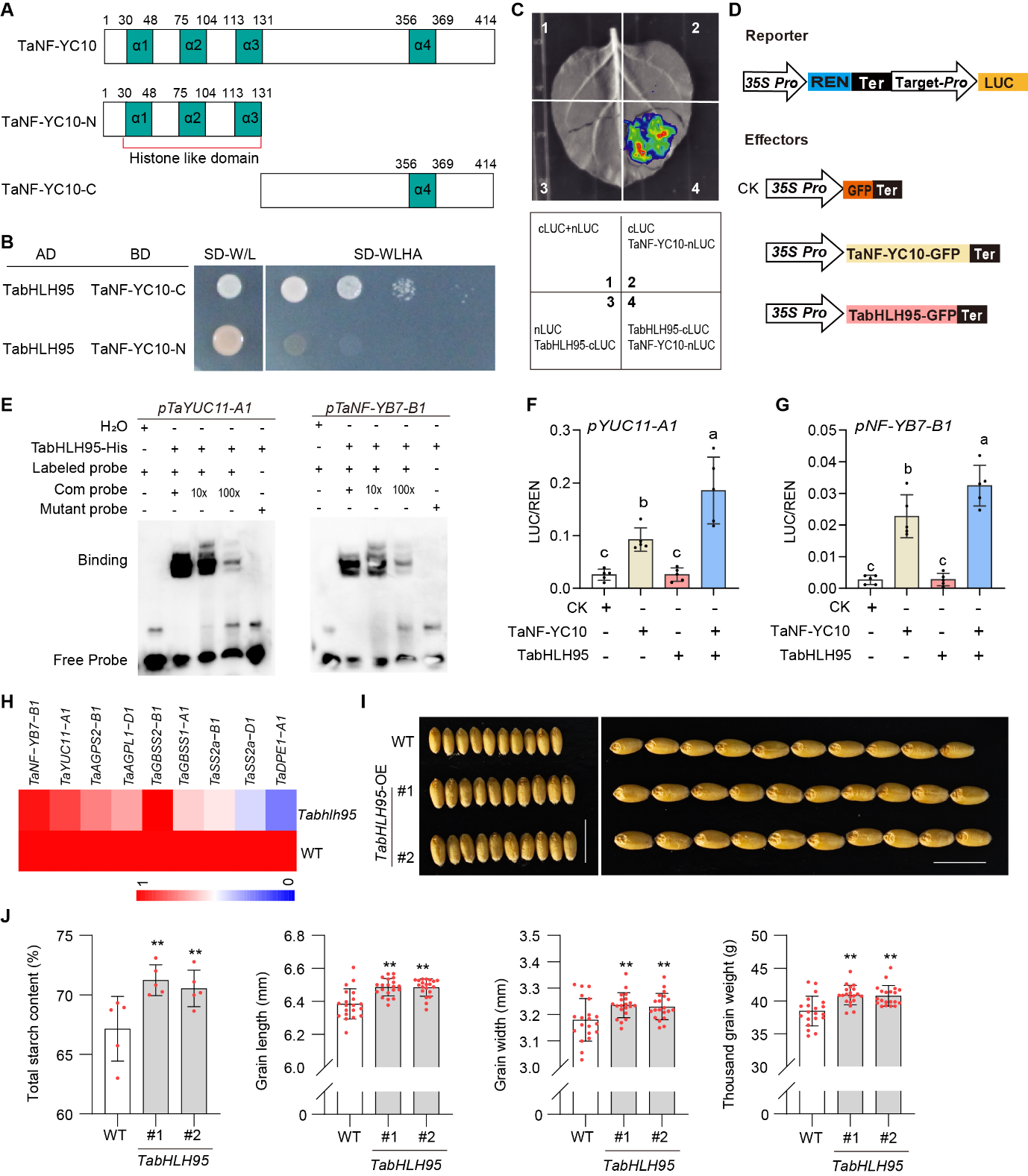

**Supplementary Figure 5. TabHLH95 interacts with TaNF-YC10 to cooperatively regulate starch biosynthesis and grain weight in wheat. (A)** Schematic diagrams of TaNF-YC10 and its truncated variants used for interaction analyses. TaNF-YC10-N harbors the N-terminal region with a histone-like domain, whereas TaNF-YC10-C contains the C-terminal α4 region. Numbers indicate amino-acid positions. **(B)** Y2H assays showing the interaction between TabHLH95 and TaNF-YC10 fragments. Yeast cells were tested for growth on SD-Trp/Leu (SD-WL) and SD-Trp/Leu/His/Ade (SD-WLHA) selective media. **(C)** LCI assays in confirming the interaction between TabHLH95 and TaNF-YC10. **(D)** Schematic representation of reporter and effector constructs used in transcriptional activation assays. **(E)** EMSAs showing that TabHLH95 binds to the promoters of *TaYUC11-A1* and *TaNF-YB7-B1*. **(F, G)** DLR assays demonstrating that TaNF-YC10 and TabHLH95 synergistically activate the promoters of *TaYUC11-A1* **(F)** and *TaNF-YB7-B1* **(G)**. Data are means ± SD (n = 5). Different lowercase letters indicate significant differences (One-way ANOVA, *P* < 0.05). **(H)** Heatmap showing relative expression levels of genes related to auxin biosynthesis and starch synthesis in WT and *Tabhlh95* mutant lines. **(I)** Phenotypes of mature grains from WT and two independent *TabHLH95*-OE lines (#1 and #2). Scale bars indicate 1 cm. **(J)** The total starch content, grain length, grain width, and thousand grain weight in WT and *TabHLH95*-OE lines. Data are means ± SD. Asterisks indicate significant differences compared with WT (One-way ANOVA, *P* < 0.01).

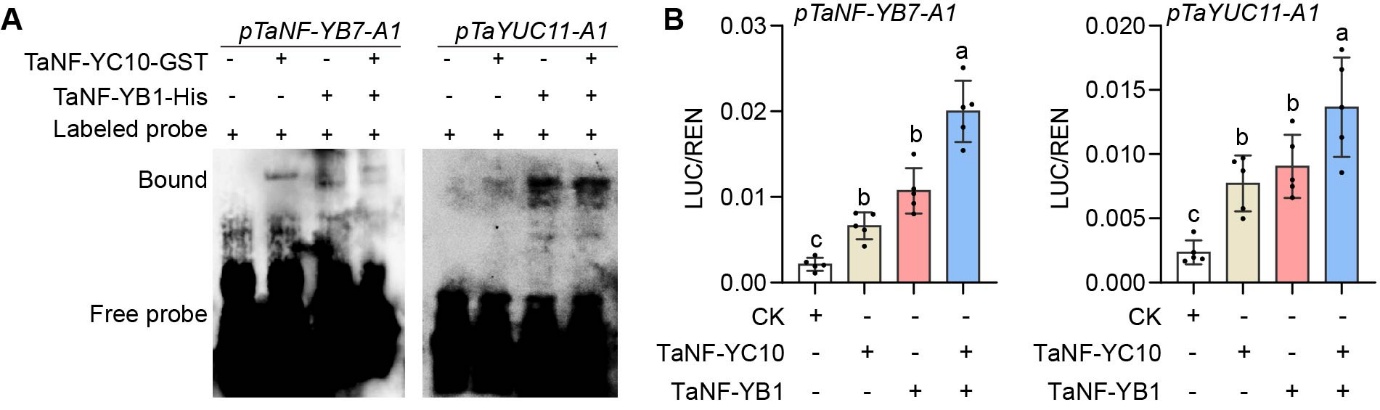

**Supplementary Figure 6. TaNF-YC10 cooperates with TaNF-YB1 to activate downstream target genes. (A)** EMSA assays showing the binding of the TaNF-YC10 and TaNF-YB1 to the promoters of *TaNF-YB7-A1* and *TaYUC11-A1*. Labelled probes corresponding to the indicated promoter regions were incubated with recombinant TaNF-YC10-GST and/or TaNF-YB-His proteins, as indicated. **(B)** DLR assays showing transcriptional activation of the *TaNF-YB7-A1* and *TaYUC11-A1* promoters by TaNF-YC10 and TaNF-YB1 in *Nicotiana benthamiana* leaves. Data are presented as means ± SD (n = 3). Statistical significance was determined by one-way ANOVA, with different letters indicating significant differences (*P* < 0.05).

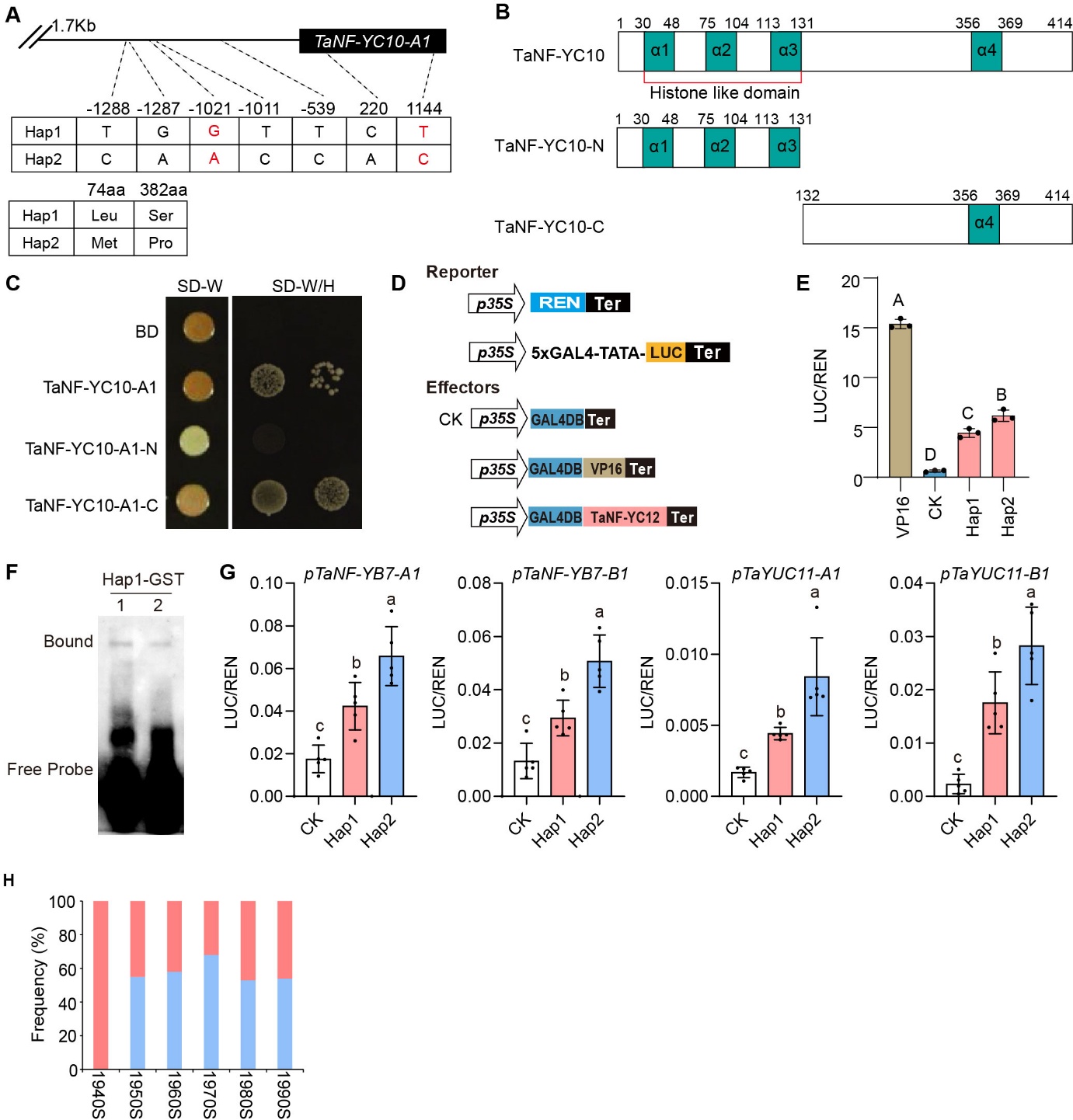

**Supplementary Figure 7. Natural variation of *TaNF-YC10-A1* leads to differential transcriptional regulation of downstream genes. (A)** Two major haplotypes of T*aNF-YC10-A1* exhibit sequence variation in both promoter and coding regions. **(B)** Domain structures of TaNF-YC10 and its truncated forms. The histone-like domain composed of α1–α3 helices is indicated, and TaNF-YC10-C contains the C-terminal α4 domain. **(C)** Yeast transcriptional activation assays showing the transactivation activity of full-length TaNF-YC10-A1 and its truncated forms. **(D)** Schematic diagrams of reporter and effector constructs used in the GAL4-based transcriptional activation assays. **(E)** DLR assays examining the transcriptional activation activities of *TaNF-YC10-A1* haplotypes. Data are presented as means ± SD (n = 3). Statistical significance was determined by one-way ANOVA, with different letters indicating significant differences (*P* < 0.05). **(F)** EMSA revealing the binding of TaNF-YC10-Hap1 to the promoter of downstream genes. 1 represents the promoter of *TaYUC11-A1* and *TaNF-YB7-A1*. **(G)** DLR assays examining the transcriptional activation capacities of *TaNF-YC10-A1* haplotypes toward downstream genes. Data are presented as means ± SD (n = 5). Statistical significance was determined by one-way ANOVA, with different letters indicating significant differences (*P* < 0.05). **(H)** Haplotype frequencies of *TaNF-YC10-A1* across different breeding years.
